# PROTEOMICS OF HYPOTHERMIC ADAPTATION REVEALS THAT RBM3 ENHANCES MITOCHONDRIAL METABOLISM AND MUSCLE STEM-CELL DIFFERENTIATION

**DOI:** 10.1101/2023.05.05.539524

**Authors:** Paulami Dey, Srujanika Rajalaxmi, Purvi Singh Thakur, Maroof Athar Hashmi, Heera Lal, Nistha Saini, Sneha Muralidharan, Raviswamy G H Math, Pushpita Saha, Swarang Sachin Pundlik, Nirpendra Singh, Arvind Ramanathan

## Abstract

Adaptation to hypothermic stress is important for skeletal muscle cells, but a comprehensive knowledge of molecular mediators is lacking. We show that adaptation to mild hypothermia (32^0^C) improves the ability of skeletal muscle myoblasts to differentiate into myotubes *in vitro*. We performed proteomic analysis of mouse myoblasts exposed to mild hypothermia for various time points and identified dynamic changes in mitochondrial metabolism and proteostasis. This revealed that RBM3, an RNA-binding protein, increases progressively with acute and chronic exposure to hypothermic stress, and is necessary for the enhanced differentiation upon hypothermic adaptation. We also demonstrate that overexpression of RBM3 at physiological temperatures is sufficient to (i) enhance mitochondrial metabolism as judged by a decrease in the AMPK energy-sensing pathway, (ii) increase levels of proteins associated with translation and increase levels of 4E-BP1 phosphorylation, (iii) increase stem cell markers (MyoD1, PAX7), and improve differentiation of myoblasts from both young and aged mice.

## Introduction

Hypothermic adaptation is a hallmark of hibernating animals, and mild hypothermia has been used for preserving tissue homeostasis in patients, including the brain and heart^1–6^. During hypothermia, mitochondrial signaling maintains cellular energy homeostasis and promotes cellular survival^7^. It has been demonstrated that exposing human myoblasts to mild hypothermic conditions can improve their transplantation efficiency and survival. This observation can be leveraged for applications of myoblasts in cell-replacement therapies^8^ and underlines the importance of uncovering comprehensive mechanisms that mediate hypothermic adaption.

In hibernating animals under hypothermic conditions (which can be as low as 0-5 ͦ C), one of the key RNA-binding proteins to be upregulated is RBM3 (RNA-binding motif protein 3) which is a cold-responsive protein^9–13^. RNA binding proteins (RBP) are characterized by the presence of an RNA binding domain (RBD) and an intrinsically disordered domain (IDD)^14^. RBPs are known to control gene expression through the regulation of protein synthesis and post-transcriptional processes like RNA splicing, RNA stability and RNA localization^15, 16^. They are also known to affect cellular physiology in the context of numerous diseases e.g. neurodegenerative, cardiovascular disorders, cancer and hyperglycemia^17–20^.

RBM3 is a highly conserved glycine-rich protein with a molecular mass of 17 kDa which is known to be involved in neuroprotection and prevention of apoptosis^21, 22^. In the context of skeletal muscles, RBM3 can inhibit atrophy in myotubes. Overexpression of RBM3 has been shown to promote survival against peroxide stress in myoblasts^23, 24^. A recent study using C2C12 cells has shown that RBM3 may bind to the 3’ UTR and coding regions of mRNA involved in various cellular processes like translational initiation and proteasomal degradation^25^.

A comprehensive knowledge of proteomic changes during hypothermia in skeletal muscle cells is lacking. Therefore, mapping the hypothermic proteome and understanding the functional role of RBM3 in cells will enable new strategies for the application of therapeutic hypothermia. To fill this gap in understanding we performed a systematic temporal analysis of hypothermic adaptation using untargeted proteomics. Our study reveals that in response to both acute and chronic hypothermia myoblasts mount a dynamic response involving mitochondrial metabolism, RNA processing and an increase in levels of RBM3. We show that overexpression of RBM3 in myoblasts at 37^0^C can recapitulate a subset of hypothermic effects on myoblast metabolism and differentiation in part by enhancing mitochondrial metabolism. Finally, we show that hypothermia and RBM3 can rejuvenate aged skeletal muscle precursor cells *in vitro* as judged by increased levels of stem cell markers (*Pax7, MyoD1*).

## Results

### Hypothermic adaptation enhances the differentiation of mouse skeletal myoblasts

To test the effects of hypothermic adaptation on skeletal muscle differentiation, we grew C2C12 myoblasts at 32^0^C and 25^0^C respectively for 72 hrs. (37^0^C was used as control). This was followed by differentiation into myotubes at 37^0^C for 6 days (Fig 1A). We observed that the mRNA levels of differentiation markers like myosin heavy chain (*MyHC*, a late differentiation marker) and myogenin (*Myog*, an early differentiation marker) increased significantly under hypothermia as compared to the control (Fig 1 B, C, D, E). The protein levels of these markers also increased significantly during hypothermia, compared to the control (Fig 1 F, 1 G). We also observed that the increase in differentiation under hypothermia was more pronounced at 25^0^C compared to that at 32^0^C (Fig S1 A). We did similar experiments using mouse primary myoblasts isolated from wild-type B6/J mice. Primary myoblasts were grown at 32^0^C for 72 hrs. and then differentiated at 37^0^C for 6 days (37^0^C was used as a control). There was a loss of viability of primary myoblasts at 25^0^C unlike C2C12 cells (data not shown). Similarly to C2C12 cells, we observed that both mRNA and protein levels of the differentiation markers significantly increased under hypothermia as compared to 37^0^C control (Fig 1 H, I, J). These results together imply that hypothermic adaptation in mouse skeletal myoblasts enhances differentiation.

**Figure 1.**
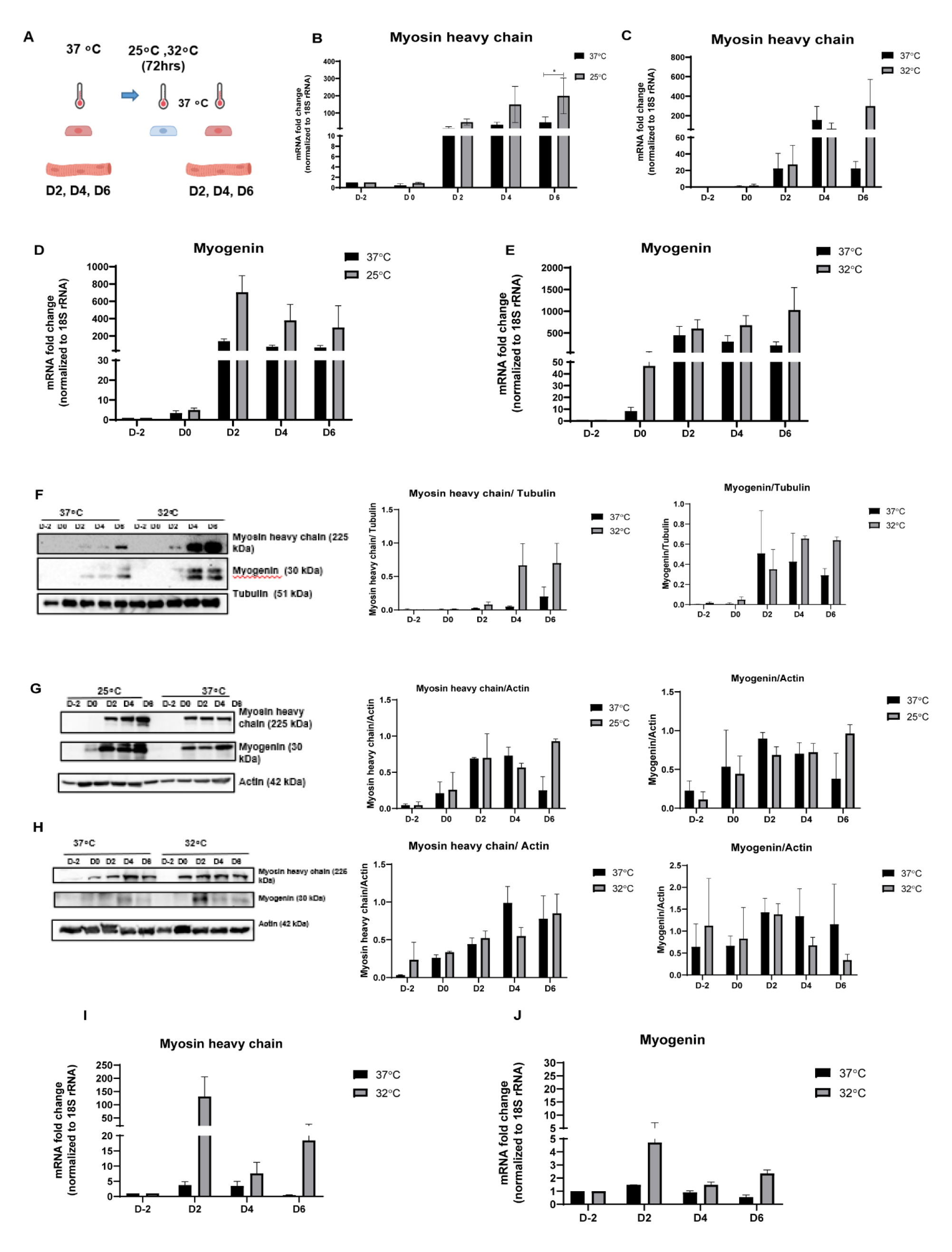
Hypothermic adaptation (after 72 hrs.) enhances differentiation of skeletal muscle cells. **(A)** Schematic representing the hypothermia experiment where C2C12 myoblasts were differentiated into myotubes at 37^0^C after 72 hrs. of hypothermia (25^0^C or 32^0^C). **(B)** mRNA expression levels of Myosin heavy chain (*MyHC*) during differentiation using C2C12 cells at 25^0^C and 37^0^C (n=3) where the x-axis represents the number of days pre-differentiation and during differentiation and the y-axis represents the mRNA fold change of *MyHC* normalized to 18S rRNA and compared to D-2. **(C)** mRNA expression levels of *MyHC* during differentiation using C2C12 cells at 32^0^C and 37^0^C (n=3) where the x-axis represents the number of days pre-differentiation and during differentiation and the y-axis represents the mRNA fold change of *MyHC* normalized to 18S rRNA and compared to D-2. **(D)** mRNA expression levels of Myogenin (*Myog*) during differentiation in C2C12 cells at 25^0^C and 37^0^C (n=3) where the x-axis represents the number of days pre-differentiation and during differentiation and the y-axis represents the mRNA fold change of *Myog* normalized to 18S rRNA and compared to D-2. **(E)** mRNA expression levels of *Myog* during differentiation using C2C12 cells at 32^0^C and 37^0^C (n=3) where the x-axis represents the number of days pre-differentiation and during differentiation and the y-axis represents the mRNA fold change of *Myog* normalized to 18S rRNA and compared to D-2. **(F)** Western blot analysis of MyHC and MYOG during differentiation using C2C12 cells at 32^0^C and 37^0^C (n=2). The bar graph represents the MyHC/Actin and MYOG/Actin levels, where the x-axis represents the number of days pre-differentiation and during differentiation and the y-axis represents relative levels of MyHC and MYOG w.r.t to Actin. **(G)** Western blot analysis of MyHC and MYOG during differentiation using C2C12 cells at 25^0^C and 37^0^C (n=1). The bar graph represents the MyHC/Actin and MYOG/Actin levels, where the x-axis represents the number of days pre-differentiation and during differentiation and the y-axis represents relative levels of MyHC and MYOG w.r.t to Actin. **(H)** Western blot analysis of MyHC and MYOG during differentiation using mouse primary myoblasts at 32^0^C and 37^0^C (n=1). **(I)** mRNA expression levels of *MyHC* during differentiation using mouse primary myoblasts cells at 32^0^C and 37^0^C where the x-axis represents the number of days pre-differentiation and during differentiation and the y-axis represents the mRNA fold change of *MyHC* normalized to 18S rRNA and compared to D-2. **(J)** mRNA expression levels of *Myog* during differentiation using mouse primary myoblasts at 32^0^C and 37^0^C where the x-axis represents the number of days pre-differentiation and during differentiation and the y-axis represents the mRNA fold change of *MyHC* normalized to 18S rRNA and compared to D-2. *, **, *** represent p-value < 0.05, 0.01 and 0.001 respectively.

### C2C12 cells grown under hypothermia (32^0^C) for 6, 12, 24 and 48 hrs. show a dynamic increase in levels of proteins involved in RNA processing and cellular metabolism

To understand underlying global changes under hypothermia, we performed proteomic profiling of C2C12 cells grown under hypothermia (32^0^C) at different time points. We grew C2C12 cells at 32^0^C for 6 hrs., 12 hrs., 24 hrs., and 48 hrs. and performed proteomic profiling using a SWATH proteomics workflow. We observed that among the 1347 proteins detected, 323 proteins were upregulated at 6 hrs. (24%), 237 proteins were upregulated at 12 hrs. (17.6%), 284 proteins were upregulated at 24 hrs. (21%) and 300 proteins were upregulated at 48 hrs. (22%) respectively. 469 proteins were downregulated at 6 hrs. (34.9%), 393 proteins were downregulated at 12 hrs. (29.2%), 413 proteins were downregulated at 24 hrs. (30.7%) and 390 proteins (29%) were downregulated at 48 hrs. respectively (Fig 3S A).

Hypothermia induced acute changes in protein levels at 6 hrs. and differential responses over 12 hrs., 24 hrs., and 48 hrs. respectively (Fig 2 A, B, C). We observed that broadly hypothermia increased proteins involved in RNA processing, lipid metabolism, tricarboxylic acid (TCA), and electron transport chain (ETC) pathways. GO pathway and process enrichment analysis also indicated that pathways involved in carbon (monocarboxylic, amino acid, and energy) metabolism and RNA processing were progressively upregulated under hypothermia (Fig 3 A, B, C, D).

**Figure 2.**
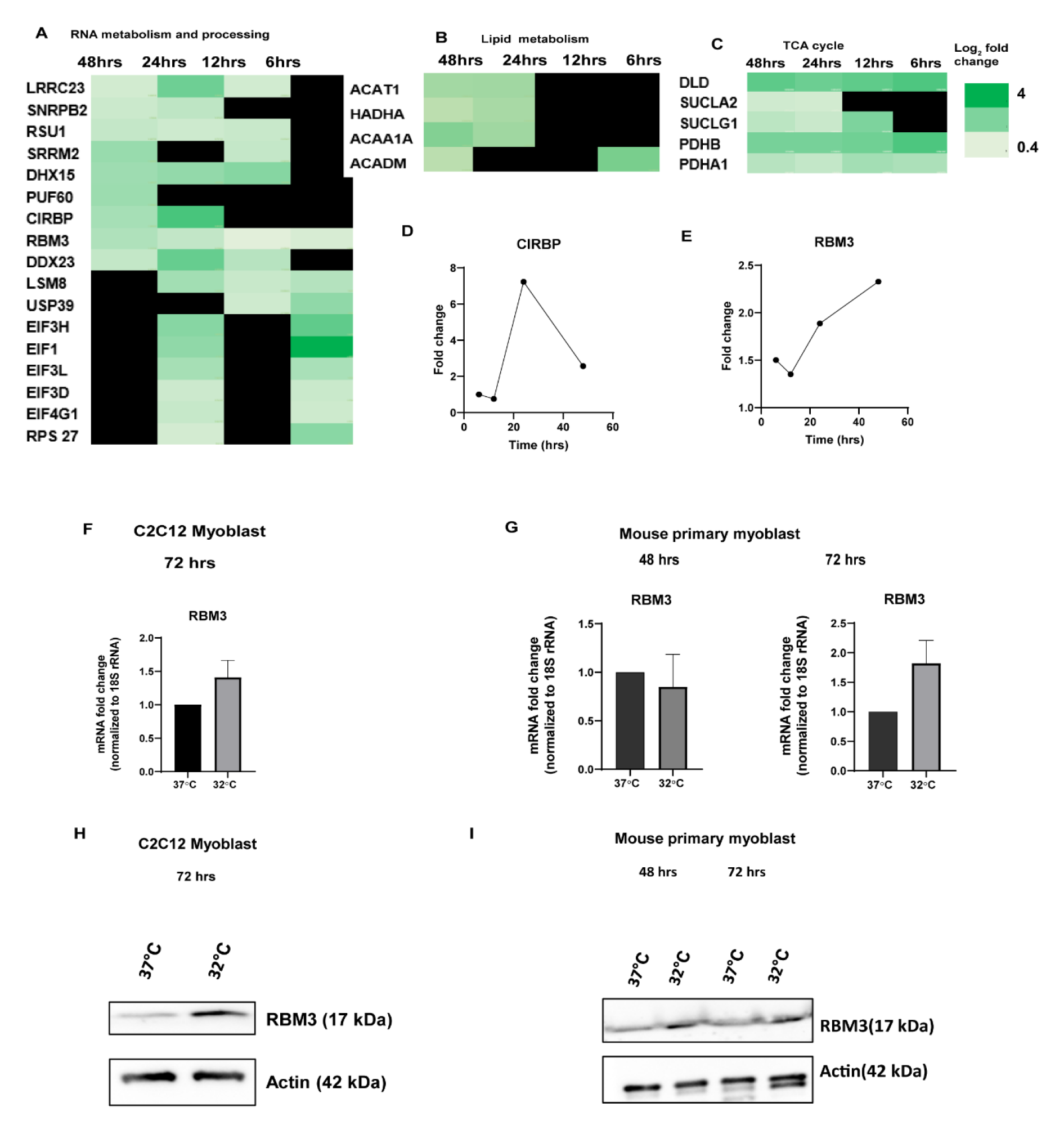
Proteomic mapping of the temporal response of C2C12 cells to hypothermia (32^0^C) at 6, 12, 24 and 48 hrs. Heat map of upregulated proteins involved in **(A)** RNA metabolism and processing. **(B)** lipid metabolism. **(C)** TCA cycle. (n=3). Graph showing fold change of protein expression of RNA binding proteins **(D)** CIRBP **(E)** RBM3 during hypothermic treatment where the x-axis represents time in hours and the y-axis represents the relative fold change of the protein in hypothermic condition compared to control. **(F)** mRNA expression levels of *RBM3* using C2C12 cells at 32^0^C and 37^0^C at 72 hrs., where the x-axis represents the temperature, and the y-axis represents the mRNA fold change of RBM3 normalized to 18S rRNA and compared to 37^0^C (n=3). **(G)** mRNA expression levels of RBM3 using mouse primary myoblasts at 32^0^C and 37^0^C at 48 hrs. and 72 hrs. respectively, where the x-axis represents the temperature and the y-axis represents the mRNA fold change of *RBM3* normalized to 18S rRNA and compared to 37^0^C (n=2). **(H)** Western blot analysis of RBM3 using C2C12 cells at 32^0^C and 37^0^C at 72 hrs. (n=2). **(I)** Western blot analysis of RBM3 using mouse primary myoblasts at 32^0^C and 37^0^C at 48 hrs. and 72 hrs. respectively (n=2).

**Figure 3.**
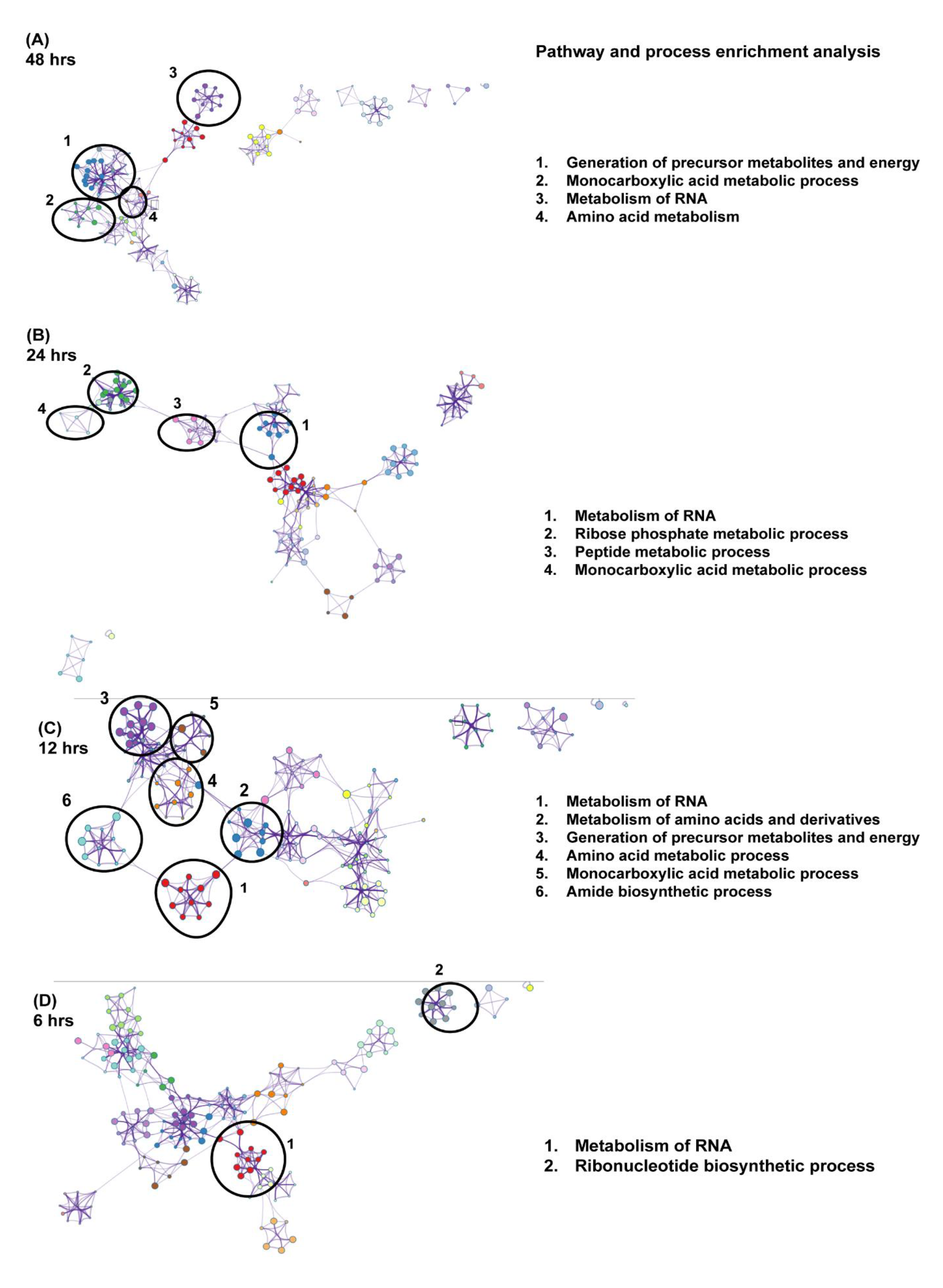
GO enrichment analysis of protein functional network of C2C12 cells in hypothermia (32^0^C) at 6, 12, 24 and 48 hrs. GO enrichment protein functional network at **(A)** 48 hrs. **(B)** 24 hrs. **(C)** 12 hrs. **(D)** 6 hrs. (n=3).

With respect to RNA metabolism and processing, at 6 hrs. there was a significant upregulation of proteins involved in translation initiation (eIF-3h, eIF-1, eIF-3i, eIF-3d, eIF-4G1, RPS27). At 48 hrs. most of these proteins were not detectable but instead, proteins involved in RNA-splicing (PUF60, DDX15, USP39, SNRPB2), and RNA processing (DHX15, LSM8) were significantly upregulated. 12 hrs. and 24 hrs. showed upregulation of proteins present within both these sets. With respect to proteins associated with lipid metabolism, a set of unique proteins involved in beta-oxidation was upregulated at 6 hrs. (Fig S2 A). A subset of these proteins involved in beta-oxidation of fatty acids (ACAT1, ACADM, HADHA, ACAA1A, HSD17B4, HSD17B10), were detected at 24 hrs. and 48 hrs. respectively. With respect to the TCA pathway, three proteins, which are components of the mitochondrial pyruvate dehydrogenase complex (DLD, PDHB, PDHA1) were upregulated at 6 hrs., 12 hrs., 24 hrs., and 48 hrs. respectively. Succinate-CoA ligase (SUCLA2, SUCLG1) was not present at 6 hrs. and 12 hrs. but upregulated significantly at 24 hrs. and 48 hrs.

Another set of unique proteins was upregulated only at 6 hrs., 12 hrs., 24 hrs., and 48 hrs. respectively and these included proteins involved in RNA splicing, RNA processing, lipid metabolism, TCA, ETC (Listed in Fig S2 A, B).

Cellular processes like RNA processing and metabolism, monocarboxylic acid metabolism, amino acid metabolism, acyl-CoA metabolism and splicing of RNA were upregulated under hypothermia at different time points as shown by the Metascape analysis software^26^ (summarized in Table 1, Fig S3 B, C, D, E).

**Table 1.** Summary of the GO enrichment pathway analysis of the cellular processes that were increased during hypothermia at different time points.

| CELLULAR PROCESSES | 6 hrs | 12 hrs | 24 hrs | 48 hrs |
| --- | --- | --- | --- | --- |
| Metabolism of RNA | + | + | + | + |
| Cellular responses to stress | + | - | + | + |
| Monocarboxylic Acid metabolic processes | - | + | + | + |
| Amino Acid Metabolism | - | + | + | + |
| Acyl coA metabolic processes | - | - | + | + |
| Ribonucleotide metabolic processes | + | - | + | - |
| Regulation of splicing | + | - | + | - |
| Carbon metabolism | - | + | - | - |

### RBM3 is upregulated by hypothermic stress in C2C12 and mouse primary myoblasts

RNA binding proteins (RBM) are a class of proteins that bind to double or single-stranded RNA. They contain an RNA-recognition motif (RRM) and are critical for RNA processing and stress response^27^. Two proteins belonging to the RBM family were upregulated in response to hypothermia: RBM3 and CIRBP (Cold inducible RNA binding protein). RBM3 was upregulated even at 6 hrs. and increased progressively till 48 hrs. whereas, CIRBP was expressed only at 24 hrs. and its level decreased by 48 hrs. (Fig 2 D, E). Therefore, we investigated the role of RBM3 in the hypothermic adaptation of myoblasts.

The mRNA levels of *RBM3* increased after 48 hrs. and 72 hrs. of hypothermia in C2C12 and mouse primary myoblasts (Fig 2 F, G).

By western blotting, we confirmed that protein levels of RBM3 increased after 48 hrs. and 72 hrs. of hypothermia in C2C12 cells and mouse primary myoblasts (Fig 2 H, I).

### RBM3 is required for hypothermia-mediated enhanced C2C12 myoblast differentiation

Since hypothermia was observed to promote differentiation of skeletal muscle myoblasts and RBM3 was upregulated during hypothermia, we hypothesized that hypothermia-dependent promotion of differentiation could be dependent on the expression of RBM3. To test this, C2C12 cells were transfected with siRBM3 (scrambled siRNA was used as control) (Fig S4 A). After 48 hrs. of siRNA treatment, we incubated the C2C12 myoblasts (siRBM3 transfected and scrambled siRNA transfected) at 32^0^C for 72 hrs. (37^0^C scrambled siRNA treated cells were used as control) followed by differentiation into myotubes at 37^0^C for 6 days. We observed that the mRNA levels of differentiation markers: *MyHC* and *Myog,* increased significantly at 32^0^C in scrambled siRNA-treated cells compared to 37^0^C scrambled siRNA-treated cells as expected. Conversely, we observed that cells treated with siRBM3 showed a decrease in differentiation markers after incubation at 32^0^C (Fig 4 A, B).

**Figure 4.**
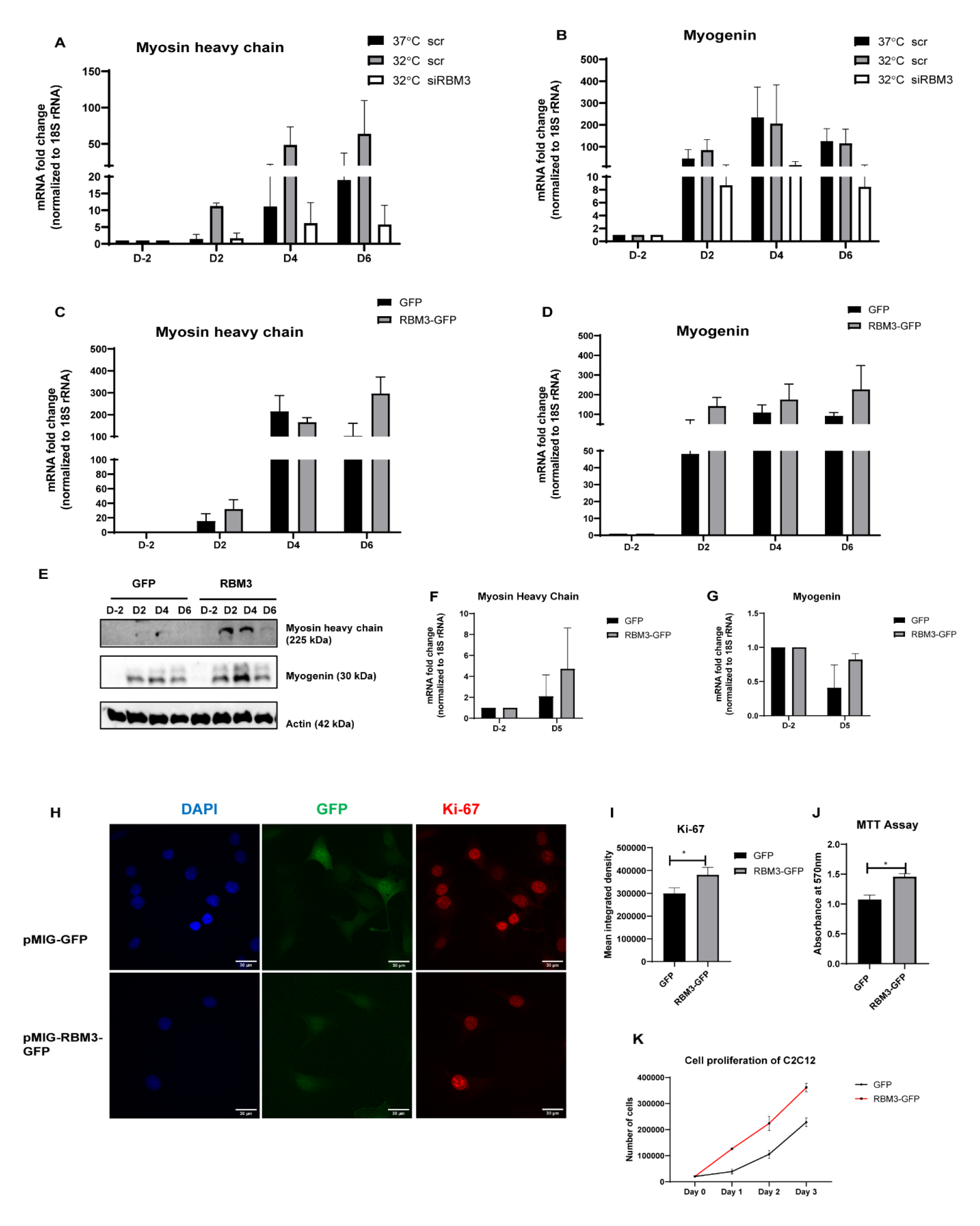
RBM3 promotes differentiation and viability of skeletal muscle myoblasts. **(A)** mRNA expression levels of *MyHC* for siRBM3 and scrambled control (scr) treatment using C2C12 cells at 32^0^C and 37^0^C (n=2), where the x-axis represents the number of days pre-differentiation and during differentiation and the y-axis represents the mRNA fold change of *MyHC* normalized to 18S rRNA and compared to D-2. **(B)** mRNA expression levels of *Myog* for siRBM3 and scrambled control (scr) treatment using C2C12 cells at 32^0^C and 37^0^C (n=2), where the x-axis represents the number of days pre-differentiation and during differentiation and the y-axis represents the mRNA fold change of *Myog* normalized to 18S rRNA and compared to D-2. **(C)** mRNA expression levels of *MyHC* during differentiation using C2C12 cells overexpressing pMIG-GFP control and pMIG-RBM3 at 37^0^C (n=3), where the x-axis represents the number of days pre-differentiation and during differentiation and the y-axis represents the mRNA fold change of *MyHC* normalized to 18S rRNA and compared to D-2. **(D)** mRNA expression levels of *Myog* during differentiation using C2C12 cells overexpressing pMIG-GFP control and pMIG-RBM3 at 37^0^C (n=3), where the x-axis represents the number of days pre-differentiation and during differentiation and the y-axis represents the mRNA fold change of *Myog* normalized to 18S rRNA and compared to D-2. **(E)** Western blot analysis of MyHC and MYOG during differentiation using C2C12 cells overexpressing pMIG-GFP control and pMIG-RBM3 at 37^0^C (n=2). **(F)** mRNA expression levels of *MyHC* during differentiation using mouse primary myoblasts overexpressing pMIG-GFP control and pMIG-RBM3 at 37^0^C (n=2), where the x-axis represents the number of days pre-differentiation and during differentiation and the y-axis represents the mRNA fold change of *MyHC* normalized to 18S rRNA and compared to D-2. **(G)** mRNA expression levels of *Myog* during differentiation using mouse primary myoblasts overexpressing pMIG-GFP control and pMIG-RBM3 at 37^0^C (n=2), where the x-axis represents the number of days pre-differentiation and during differentiation and the y-axis represents the mRNA fold change of *Myog* normalized to 18S rRNA and compared to D-2. **(H)** Confocal images of C2C12 cells overexpressing pMIG-GFP control and pMIG-RBM3 at 37^0^C. Red indicates Ki-67 staining, blue indicates DAPI, and Green indicate GFP (Scale bar, 30um). **(I)** Bar graph representing the quantitation of Ki-67 staining where the y-axis represents the mean integrated density of Ki-67 positive nucleus stain. **(J)** Bar graph for MTT assay using C2C12 cells overexpressing pMIG-GFP control and pMIG-RBM3 at 37^0^C where the Y axis represents absorbance at 570 nm (n=3). **(K)** Line graph indicating cell proliferation of C2C12 cells overexpressing pMIG-GFP control and pMIG-RBM3 at 37^0^C where the Y-axis represents the total number of cells and X-axis represents time in days (n=3). *, **, *** represents p-value < 0.05, 0.01 and 0.001 respectively.

### Overexpression of RBM3 is sufficient to enhance differentiation of C2C12 and mouse primary myoblast at 37^0^C

Since RBM3 was required for hypothermia-driven enhanced differentiation of myoblasts, we tested whether RBM3 was sufficient for enhanced differentiation of myoblasts at 37^0^C. We generated stable C2C12 cell lines overexpressing pMIG-RBM3 (pMIG-GFP was used as control) (Fig S4 B, C). Stable C2C12 cells (overexpressing RBM3) were grown at 37^0^C and differentiated at 37^0^C for 6 days. We observed that mRNA and protein levels of the differentiation marker MyHC increased significantly in C2C12 cells overexpressing RBM3 compared to the control. This indicates that RBM3 can promote the differentiation of myoblasts in a hypothermia-independent manner (Fig 4 C, D).

We performed a similar experiment in mouse primary myoblasts where we transfected the primary cells with pMIG-RBM3 (pMIG-GFP was used as control) and differentiated them at 37^0^C for 6 days. We observed similar results of increased mRNA levels of differentiation markers (*MyHC and Myog*) due to overexpression of RBM3. (Fig 4 F, G)

### Overexpression of RBM3 promotes C2C12 myoblasts viability and proliferation

Next, we tested the effects of RBM3 on cell proliferation and cell viability of mouse skeletal myoblasts. We grew the stable lines of C2C12 cells on coverslips and stained them with Ki-67 antibody and performed confocal microscopy to determine the mean integrated intensity of the Ki-67 positive nuclei. We observed that overexpression of RBM3 leads to an increase in the Ki- 67 intensity. (Fig 4 H, I). We also performed a cell counting assay where we plated an equal number of cells (of both GFP and RBM3) respectively and counted the cell number from day 1 to day 3. We observed that overexpression of RBM3 leads to an increase in cell proliferation in all three days (Fig 4 K). Next, we tested whether the overexpression of RBM3 affected cell viability. We grew the C2C12 cells in 96 well plates for two days and performed an MTT assay. We observed that overexpression of RBM3 increased cell viability compared to the control (Fig 4 J). We also checked for the levels of the myoblast marker *MyoD1* in mouse primary cells transfected with RBM3 (GFP-transfected cells were used as control). We observed that mRNA expression levels of *MyoD1* increased with overexpression of RBM3 compared to control cells (Fig S4 D). These observations suggest that overexpression of RBM3 promoted cell viability, proliferation of C2C12 myoblasts and increased expression levels of the undifferentiated myoblast marker.

### Overexpression of RBM3 in C2C12 myoblasts upregulates subsets of processes involved in hypothermia, specifically those involved in RNA processing, lipid, and mitochondrial metabolism

Using proteomics, we observed that the overexpression of RBM3 significantly upregulated levels of proteins involved in RNA metabolism and processing: for example, proteins involved in proteasomal degradation (PSMC4, PSME2, PSMD14, PSMD6, PSMD8) and RNA splicing (SNRNP200, SF3B5, SRRM2). We found proteins involved in lipid metabolism (ACADL, ACADSB, ME2, ECHS1, FABP5) to be upregulated. Proteins involved in ETC (NDUFA5, NDUFA2, NDUFV1) were also upregulated (Fig 5 A, B, C). Cellular processes like RNA metabolism, fatty acid metabolism, ribonucleoprotein complex biogenesis, and RNA localization were upregulated significantly as analyzed by Metascape software (Fig 5 E). GO pathway and process enrichment analysis revealed that pathways involved in the RNA metabolism, RNA localization, and fatty acid metabolism were upregulated due to overexpression of RBM3 (Fig S5 A). We compared the hypothermia-responsive proteome and the RBM3 overexpression-responsive proteome. Of the 300 proteins that were upregulated during chronic exposure to hypothermia (48 hrs.) and 302 upregulated proteins during overexpression of RBM3, seventy-seven proteins were upregulated under both these conditions. Of this subset, several proteins were involved in lipid and mitochondrial pathways (Fig 5 D). Based on these observations we performed a lipidomic analysis of C2C12 overexpressing pMIG-RBM3 (pMIG-GFP was used as control). We observed that overexpression of RBM3 decreased the levels of triglycerides (TG) and cholesterol esters compared to the control (S9 A, B). This revealed a systemic rewiring of the metabolism of myoblasts by RBM3.

**Figure 5.**
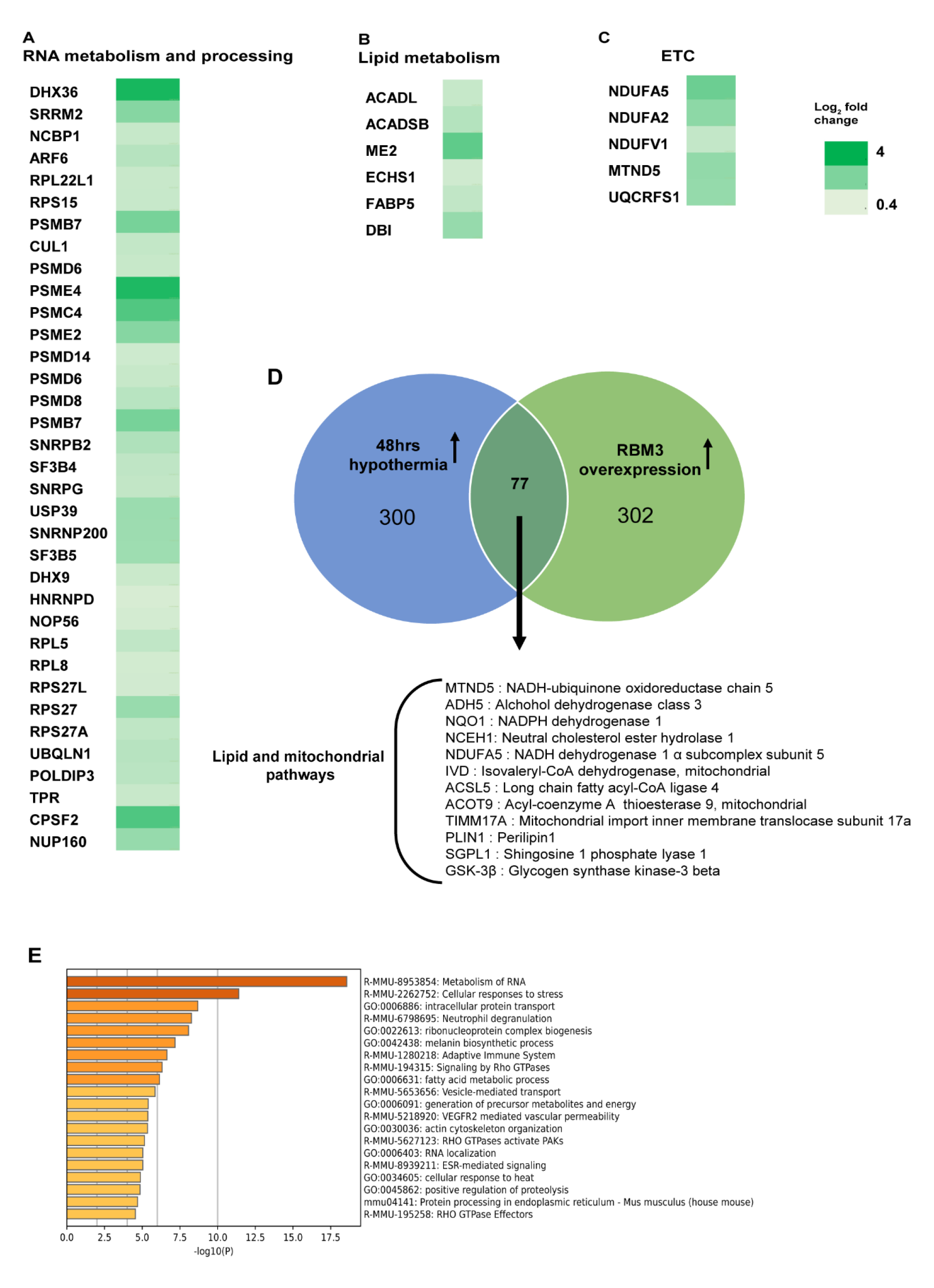
Proteomic mapping of C2C12 cells overexpressing RBM3. Heat map of upregulated proteins involved in **(A)** RNA metabolism and processing. **(B)** lipid metabolism. **(C)** ETC. **(D)** Venn diagram representing the upregulated proteins in 48 hrs. hypothermic treatment and RBM3 overexpression and the common proteins between them in C2C12 cells. **(E)** GO enrichment pathway analysis of upregulated proteins in RBM3 overexpression (n=3).

### Overexpression of RBM3 promotes mitochondrial metabolism, levels of intracellular Acetyl-CoA, and levels of the PKM2 splice variant

Since our proteomic studies revealed that overexpression of RBM3 increased the lipid and mitochondrial metabolic processes, we next investigated levels of associated metabolic pathways.

We performed a seahorse analysis of stable C2C12 cells-overexpressing RBM3 (pMIG-RBM3) (pMIG-GFP was used as control) to measure oxygen consumption rate (OCR) of the cells, under basal conditions and in the presence of metabolic inhibitors (Fig 6 A). Overexpression of RBM3 increased the basal respiration of C2C12 myoblasts compared to the control. The maximum respiration, spare respiratory capacity, and ATP-linked respiration increased in cells overexpressing RBM3 compared to the control (Fig 6 B). This suggests that RBM3 mediates an improvement in the oxygen consumption rate.

**Figure 6.**
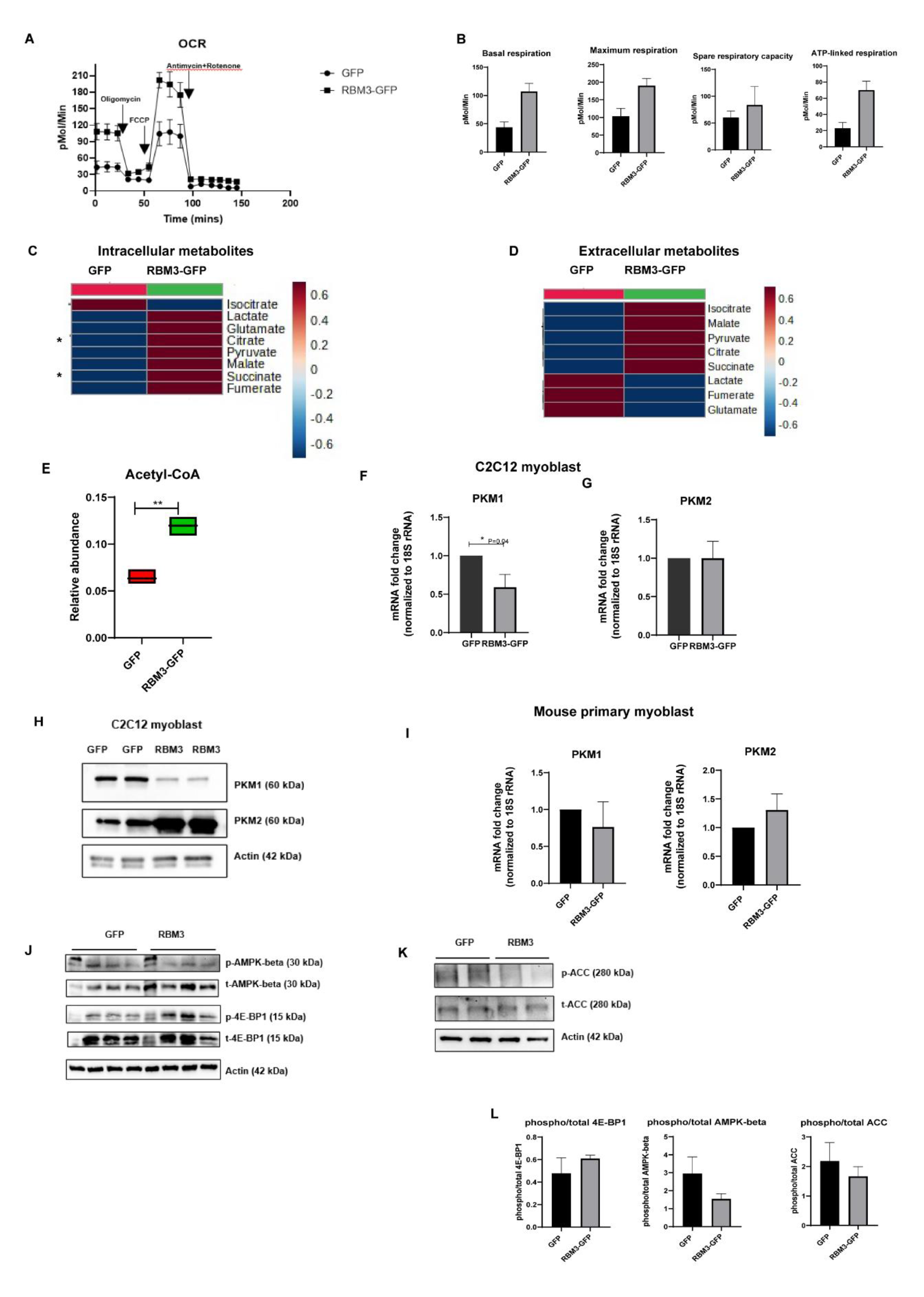
RBM3 promotes mitochondrial metabolism of skeletal muscle myoblasts. **(A)** Graph showing the mitochondrial OCR of C2C12 cells overexpressing pMIG GFP and pMIG RBM3-GFP, basal OCR and OCR after treatment with oligomycin (1 uM), FCCP (3 uM), antimycin and rotenone (1.5 uM), where the x-axis represents time in minutes and the y-axis represents oxygen consumption rate in pMol/min. **(B)** Bar graph measuring the basal respiration, maximum respiration (OCR after FCCP addition), spare respiratory capacity (basal respiration-maximum respiration) and ATP-linked respiration (basal respiration-respiration after oligomycin addition) of C2C12 cells overexpressing pMIG-GFP and pMIG-RBM3 GFP where the y-axis represents oxygen consumption rate in pMol/Min (n=2). **(C)** Heat map showing levels of TCA metabolites using C2C12 cells overexpressing pMIG-GFP control and pMIG-RBM3. **(D)** Heat map showing levels of TCA metabolites using media (48 hrs.) from C2C12 cells overexpressing pMIG-GFP control and pMIG-RBM3 (n=3). **(E)** Graphical representation of levels of acetyl-CoA using C2C12 cells overexpressing pMIG-GFP control and pMIG-RBM3 (n=3). mRNA expression levels of glycolytic genes **(F)** *PKM1*, **(G)** *PKM2* in C2C12 overexpressing pMIG-GFP control and pMIG-RBM3. **(H)** Western blot analysis of glycolytic protein levels (PKM1, PKM2) using C2C12 cells overexpressing pMIG-GFP control and pMIG-RBM3. **(I)** mRNA expression levels of glycolytic genes *PKM1*, *PKM2* in mouse primary myoblasts overexpressing pMIG-GFP control and pMIG-RBM3 (n=3). **(J)** Western blot analysis of AMPK-beta and 4E-BP1 using C2C12 cells overexpressing pMIG-GFP control and pMIG-RBM3. **(K)** Western blot analysis of acetyl-CoA carboxylase (ACC) using C2C12 cells overexpressing pMIG-GFP control and pMIG-RBM3. **(L)** Bar graph quantifying phosphorylated/total 4E-BP1, ACC and AMPK-beta respectively. *, **, *** represents p-value < 0.05, 0.01 and 0.001 respectively.

We performed an MTT assay to test the viability of cells overexpressing RBM3 in the presence of mitochondrial and glycolytic inhibitors. We observed a decreased sensitivity of cells overexpressing RBM3 to oligomycin (mitochondrial complex V inhibitor) and 2DG (2- deoxyglucose, a competitive inhibitor of glucose) compared to the control (Fig S6 A, C), as judged by the increase in the IC50 value of the inhibitors in cells overexpressing RBM3 compared to the control (Fig S6 B, D).

Since we observed an increase in mitochondrial oxygen consumption rate in cells overexpressing RBM3, we tested if there is any difference in mitochondrial morphology or mitochondrial membrane potential using mitotracker and TMRM staining respectively. We observed no significant differences in mitochondrial morphology and membrane potential due to the overexpression of RBM3 (Fig S7 A, B).

We measured both intracellular and extracellular levels of glycolytic and TCA intermediates. We observed that intracellular levels of pyruvate, lactate, malate, succinate, glutamate, and fumarate increase in cells overexpressing RBM3 compared to the control. Whereas extracellular levels of lactate, glutamate, and fumarate decrease in cells overexpressing RBM3, and levels of malate, succinate and pyruvate, citrate, and isocitrate increase due to overexpression of RBM3 (Fig 6 C, D). Intracellular lactate/pyruvate ratio decreased in cells overexpressing RBM3 compared to the control (Fig S7 F, G). Intracellular acetyl-CoA levels were significantly increased in cells overexpressing RBM3 compared to the control (Fig 6 E). Concomitantly we observed that overexpression of RBM3 was associated with an increase in protein levels of pyruvate dehydrogenase (PDH) (Fig S7 C, D) and pyruvate kinase isoform 2 (PKM2) (Fig 6 F, G, H) whereas there was a decrease in succinate dehydrogenase (SDHA, SDHC) (Fig S7 C, D) and pyruvate kinase isoform1 (PKM1) (Fig 6 F, G, H). We measured the expression levels of mRNA of *Pkm1,Pkm2, Sdha, Sdhc, Pdhb* in mouse primary myoblasts and observed a decrease in *Pkm1, Sdha, Sdhb, Pdhb* and an increase in *Pkm2* (Fig 6I, S7 E).

### Overexpression of RBM3 (i) decreases phosphorylation of AMPK-beta (Ser 182), (ii) decreases phosphorylation of acetyl-CoA carboxylase (ACC) (Ser 79) and (iii) increases phosphorylation of 4E-BP1 (Thr 37/46)

RBM3 increased mitochondrial metabolism (Fig 6) and one of the key sensors of mitochondrial bioenergetics is AMP-activated protein kinase (AMPK)^27–29^. Therefore, we analysed the levels of phosphorylation of AMPK subunits and its direct phosphorylation target (ACC)^30, 31^. Western-blot analysis of C2C12 cells overexpressing RBM3 (using GFP as control) showed a decrease in the levels of phosphorylation of AMPK-beta (Ser 182) (Fig 6J, K, L). We observed a decrease in levels of phosphorylation of ACC (Ser 79) (Fig 6J, K, L). We did not observe any significant change in the ratio of phosphorylated to total AMPK-alpha (Fig S7 H, I, J).

Overexpression of RBM3 increased the levels of proteins involved in translation initiation (Fig 5 A) and phosphorylation of 4E-BP1 is known to promote cap-dependent translation. Phosphorylation of 4E-BP1 (Thr 37/46) is mediated by mTOR and it primes 4E-BP1 for getting phosphorylated at Ser 65/70 which can subsequently lead to activation of cap-dependent translation^32^. Therefore, we measured the levels of phosphorylated 4E-BP1. We observed an increase in the levels of phosphorylation of 4E-BP1 (Thr 37/46) with overexpression of RBM3 (Fig 6 J, K, L).

### Myoblasts from aged mice have a diminished ability to differentiate *in vitro* which is partially rescued by hypothermia

We evaluated whether hypothermic adaptation could improve the differentiation of aged mouse myoblasts. We isolated mouse myoblasts from aged (18 months old) and young (2-3 months old) wild-type mice (B6/J) and cultured them at 37^0^C followed by differentiation at 37^0^C for 5 days. A decrease in *MyHC* and *Myog* mRNA levels was observed in the case of aged cells as compared to the young ones (Fig S10 A, B). Additionally, we observed a decrease in the mRNA levels of stem cell marker (*Pax7*) and myoblast markers like *Myf5* and *MyoD1* in aged cells compared to young cells (Fig 7A, B, C). We next cultured the aged myoblasts in hypothermia (32^0^C for 72 hrs.) followed by differentiation at 37^0^C for 5 days (37^0^C was used as control). We observed that hypothermia increased the levels of *MyHC* (Fig 7 D) and *Myog* (Fig S10 C) in aged cells compared to the control. Also, the mRNA levels of *RBM3* increased as expected after hypothermia in aged myoblasts (Fig 7 E).

**Figure 7:**
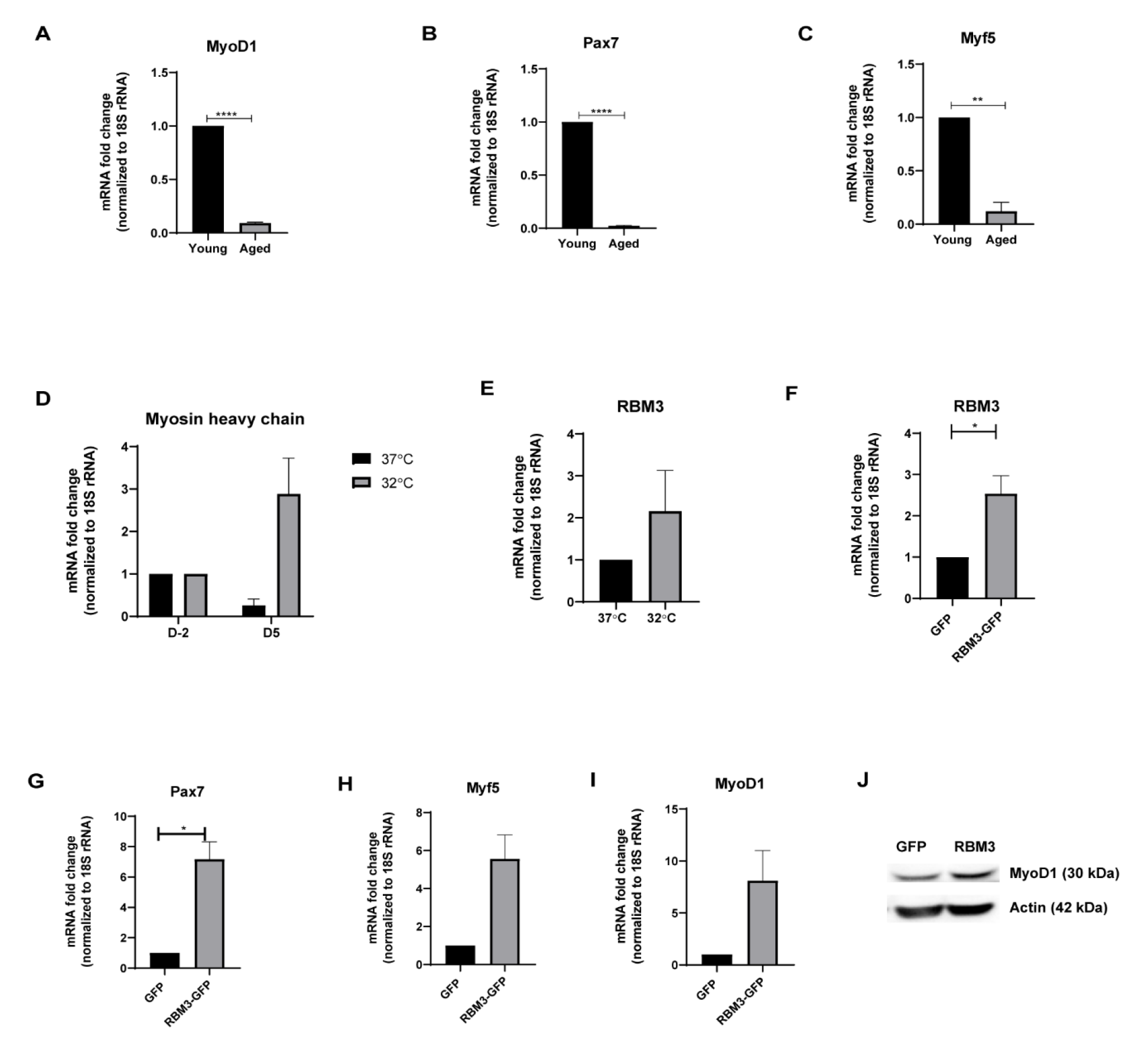
Overexpression of RBM3 increases stem cell markers in aged myoblasts. mRNA expression levels of **(A)** *MyoD1* **(B)** *Pax7* **(C)** *Myf5* in aged myoblasts compared to young **(D)** mRNA expression levels of *MyHC* in aged myoblasts at 32^0^C compared to 37^0^C. (n=2) **(E)** mRNA expression levels of *RBM3* in aged myoblasts at 32^0^C compared to 37^0^C. mRNA expression levels of **(F)** *RBM3*, **(G)** *Pax7*, **(H)** *Myf5*, **(I)** *MyoD1* in aged myoblasts transfected with RBM3 and control GFP. (n=2 for all the mRNA studies) **(J)** Western blot analysis of MyoD1 using mouse primary myoblasts at proliferative stage (72 hrs. post-transfection) overexpressing RBM3-GFP and GFP control (n=2). *, **, *** represents p-value < 0.05, 0.01 and 0.001 respectively.

**Figure 8:**
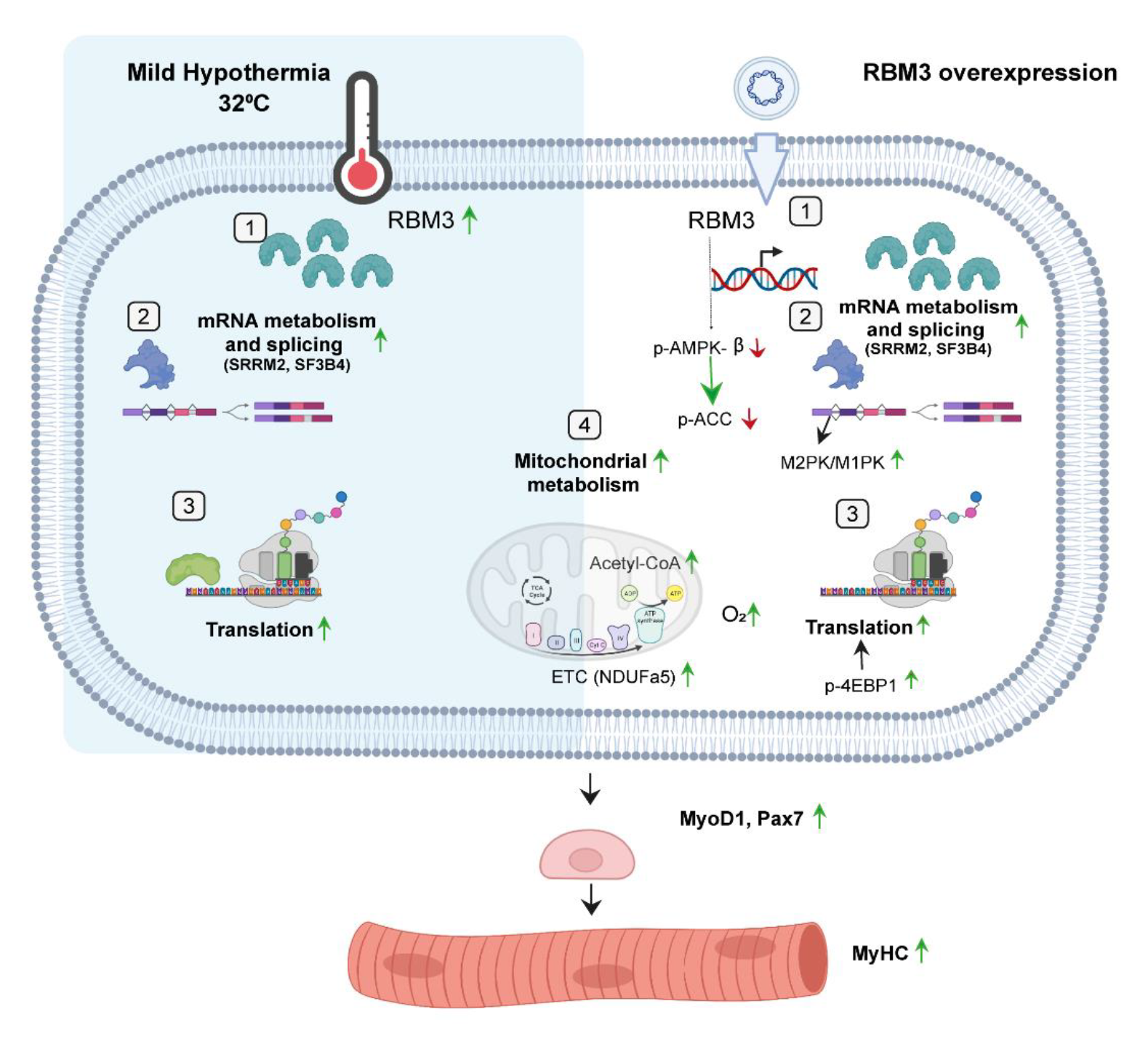
Graphical representation showing the effect of mild hypothermia and RBM3 overexpression on important cellular processes like RBM3 expression (1), RNA metabolism and splicing (2), translation (3) and mitochondrial metabolism (4) which finally leads to improved skeletal muscle regeneration.

### Overexpression of RBM3 increases mRNA and protein expression of myoblast markers in aged cells

Next, we tested whether overexpression of RBM3 was sufficient to increase the expression of stemness markers and enhance the differentiation of aged myoblasts. We transfected aged myoblasts with pMIG-RBM3 (pMIG-GFP was used as control). We observed that overexpression of RBM3 in aged cells increased the mRNA expression levels of the stem cell marker, *Pax7* and myoblast markers, *MyoD1* and *Myf5* and proteins levels of MyoD1 compared to the control (Fig 7G, H, I). We measured extracellular levels of lactate, pyruvate and TCA cycle intermediates (Malate, Succinate, Fumarate). We observed that at 24 and 48 hrs., there was a time-dependent increase in extracellular levels of TCA cycle intermediates in RBM3 overexpressing cells although these were not statistically significant (Fig S10 E, F).

## Discussion

Therapeutic hypothermia is used for treatment in patients when there is a need for preserving tissue homeostasis in cases of brain or cardiac injury^1, 2, 4, 5^. Hypothermia is also known to improve the transplantation efficiencies of human satellite cells^21^. Although this is a well-known phenomenon, the molecular players in the context of skeletal muscle remain poorly understood.

Uncovering these molecular players has key implications for new therapeutic targets in preserving or promoting skeletal muscle homeostasis. In agreement with previous observations, we found in our study that hypothermic adaptation in both C2C12 and primary myoblasts enhanced the ability of myoblasts to differentiate as judged by the mRNA-levels of differentiation markers. To understand the dynamics and dissect pathways and mechanisms underlying hypothermic adaptation we grew C2C12 cells at 32^0^C for multiple time points (6, 12, 24, and 48 hrs.) and performed unbiased proteomic profiling of intracellular proteins. We discovered a proteome-wide rewiring in response to both acute (6 hrs.) and chronic (48 hrs.) hypothermia in C2C12 myoblasts. Interestingly we observed that out of 1347 detectable proteins, 25% were upregulated as early as 6 hrs., and this reduced to approximately 21% at 48 hrs. (Fig S3 A). Concomitantly we observed a subset of translation initiation factors (eIF-3d, eIF-3i, eIF- 1) and proteins associated with the proteasomal machinery (PSMD1, PSMD3, PSME4, PSMB4) to be upregulated as early as 6 hrs., indicating that these pathways play critical in rewiring the proteome during hypothermia (Fig 2 A, Fig S2 B). We also observe a subset of proteins involved in splicing and RNA metabolism to be upregulated during hypothermia suggesting that RNA metabolism and splicing play also play an important role. A significant part of the proteome was downregulated during hypothermia. 34.9% of proteins were downregulated at an early time-point of 6 hrs. which decreased to 29% at 48 hrs. (Fig S3 A). While a subset of translation factors increased during hypothermia, there was a subset of translation factors that were also downregulated during hypothermia (eIF-3A, eIF-3F, eIF-3B) (data not shown) suggesting a context-dependent role for these factors. Apart from RNA metabolism, some of the proteins involved in mitochondrial metabolism; TCA (SUCLA2, SUCLG1), ETC (NDUFS, COX, ATP5), and lipid metabolism (ACADM, ACADSB) increased in levels during hypothermia (Fig 2 A, Fig S2 A).

Twp RNA-binding proteins increased in levels during hypothermia: RBM3 (RNA binding motif protein 3) and CIRBP (Cold inducible RNA-Binding protein). RBMs (RNA binding proteins) are a class of proteins known to be involved in the regulation of RNA. RBM3 was seen to be upregulated from early time points of 6 hrs. and gradually increased till 48 hrs. but CIRBP was downregulated during early time points but upregulated at later time points of 24 hrs. and 48 hrs. (Fig 2 D, E). Few other RNA-binding proteins like RBM25, RBM39, and RO60 were significantly downregulated.

RBM3 is a 17 kDa protein that consists of an RNA-binding domain (RBD) and an (IDD)^14^. It has been previously shown that RBM3 is neuroprotective and promotes cellular proliferation in neurons^22, 23^. It has also been previously shown that RBM3 can protect myoblasts against ROS stress^24^. A comprehensive measurement of molecular changes that mediate the effect of hypothermia and RBM3, especially in the context of skeletal muscle function have remained less understood. In this study, we have shown that hypothermia-mediated enhancement in skeletal muscle differentiation is dependent on RBM3 expression (Fig 4 A, B). Furthermore, we observed that overexpression of RBM3 is sufficient to promote skeletal muscle differentiation, independent of hypothermia (Fig 4 C, D, E) We also find that overexpression of RBM3 increases cell growth rate increases cell viability, and enhances the levels of stemness markers like and MyoD1 in skeletal muscle myoblasts (Fig 4 H, I, J, K, Fig S4 C). We performed a proteomics analysis of C2C12 cells overexpressing RBM3 to understand cellular processes that are under its control. We observed increased levels of proteins involved in processes associated with RNA metabolism, similar to that observed in hypothermia (Fig 2 A, 5 A). These included proteins involved in splicing (SF3B5, SRRM2), proteasomal degradation machinery (PSMD6, PSME4, PSMC4, PSMD8). We also observed that the ratio of PKM2 to PKM1 proteins is increased in RBM3 overexpressing cells. PKM1 and PKM2 are splice variants of the glycolytic enzyme pyruvate kinase, and increased PKM2 is a marker of a stem cell state^33^. The increased PKM2 levels in RBM3 overexpression indicate an increased proliferation of cells overexpressing RBM3 compared to the control (Fig 4H, K). These results indicate that RBM3 is involved in the regulation of splicing in myoblasts.

We also observed an increase in levels of ribosomal proteins (RPS27A, RPL5, RPS27) involved in translation from the proteomic analysis (Fig 5 A). It is known that 4E-BP1 activates cap-dependent translation. mTOR mediates phosphorylation of 4E-BP1 (Thr 37/46) and it allows 4E-BP1 to get phosphorylated at Ser 65/70 which can subsequently lead to activation of cap-dependent translation^32^. Therefore, we hypothesized that overexpression of RBM3 can affect the phosphorylation of 4E-BP1 (Thr 37/46) thereby affecting translation. In this study, we observed an increase in the levels of phosphorylation of 4E-BP1 (Thr 37/46) with overexpression of RBM3. This suggests an activation of cap-dependent translation in cells overexpressing RBM3 (Fig 6J, K, L). Proteomic analysis of myoblasts overexpressing RBM3 suggests an increase in the levels of proteins involved in mitochondrial metabolism including lipid metabolism (ACADL, ACASB) and ETC (NDUFS) (Fig 5 B, C).

Overall, this shows that RBM3 can recapitulate aspects of hypothermia even at room temperature.

As RBM3 upregulated levels of proteins involved in lipid metabolism, like beta-oxidation of fatty acids, we performed lipidomics analysis of RBM3 overexpressed cells, We found that the levels of triglycerides and cholesterol esters were decreased in cells overexpressing RBM3 (Fig S9).

This suggests that RBM3 may cause increased mobilization of fatty acids which correlates with decreased levels of triglycerides. The decrease in cholesterol esters may imply the role of RBM3 in cholesterol metabolism as well. In this context, the increased intracellular levels of acetyl-CoA could be indicative of enhanced beta-oxidation of lipids (Fig 6 E).

In order to gain a deeper perspective of the role of RBM3 in regulating metabolism, we analyzed oxygen consumption using a Seahorse instrument. We showed that overexpression of RBM3 improved basal respiration. The maximum respiration of the cells after treatment with the uncoupler FCCP also increased in the presence of RBM3 (Fig 6 A, B). This could suggest an increase in the levels of proteins involved in the electron transport chain. In agreement with this, from the proteomics analysis, we observed an increase in mitochondrial ETC proteins like the complex I proteins NDUF1, NDUF3 and NDUFV1 (Fig 5 C). The TMRM and mitotracker imaging experiment revealed no significant change in mitochondrial membrane potential and mitochondrial morphology (Fig S7 A, B). This suggests that an increase in maximal respiration could be due to an increase in the levels of proteins involved in the mitochondrial electron transport chain, and not the overall mitochondrial biogenesis. To further verify these results, we performed metabolomics analysis of intracellular and extracellular TCA and glycolytic metabolites. We observed that the intracellular levels of citrate, pyruvate, succinate, and malate increased overexpressing RBM3. Concomitantly extracellular levels of citrate and isocitrate were increased, supporting an enhanced mitochondrial metabolism upon overexpression of RBM3 (Fig 6 C, D).

We observed an increase in mitochondrial oxygen consumption rate (OCR) and ATP-linked respiration in cells overexpressing RBM3 (Fig 6A, B) which shows increased mitochondrial metabolism. It is known that AMPK plays a vital role in regulating mitochondrial energy homeostasis by sensing the levels of intracellular AMP^8^. AMPK is a heterotrimeric enzyme complex containing alpha, beta and gamma subunits^29–31^. Under conditions of energy-stress (high AMP to ATP ratio) in the cell, AMPK undergoes phosphorylation including the beta subunit (Ser 182)^34^. Upon phosphorylation, AMPK phosphorylates its canonical substrate ACC (Ser 79) and enhances catabolic processes in cells^29–31^. We hypothesized that an increase in mitochondrial metabolism by RBM3 could decrease the phosphorylation of AMPK. In agreement with this, we observed that overexpression of RBM3 decreases the level of phosphorylation of AMPK-beta (Ser 182) and its canonical substrate ACC (ser 79). Overall, this suggests improved energy homeostasis in cells overexpressing RBM3 (Fig 6J, K, L).

It is known that mitochondrial oxidative phosphorylation and bioenergetics play an important role in the ageing of muscle stem cells^35^. AMPK is a key node regulating mitochondrial metabolism and has been implicated in ageing^36–38^. Since we have shown that RBM3 enhances mitochondrial metabolism and affects AMPK phosphorylation, therefore we tested the effect of RBM3 in the rejuvenation of aged satellite cells. It has been shown that satellite cells from aged muscles have diminished regenerative capacity both due to cell-autonomous and non-autonomous factors^39^. RBM3 is highly expressed in long-lived strains of mice and has been shown to increase the survival of C2C12 cells under ROS and ER stress^24, 40^. We explored the causal role of RBM3 in diminishing markers of ageing in satellite cells isolated from aged mice. We show that overexpression of RBM3 can rejuvenate ageing skeletal muscle myoblasts as judged by the increased expression levels of the canonical satellite cell marker (*Pax7*) and myoblasts markers (*MyoD1* and *Myf5*) (Fig 7G, H, I, J). Moreover, we also show that overexpression of RBM3 increases the ability of aged myoblasts to produce higher levels of extra-cellular TCA intermediates e.g., succinate, malate and fumerate (Fig S10 G, H). Overall, this suggests that RBM3 may enhance satellite cell markers and metabolism *in vitro*. The study shows that RBM3 may intersect with pathways important for rejuvenating myoblasts from both young and old mice. It also provides an important underlying mechanism by which the hypothermic cells control multiple aspects of cellular physiology like RNA metabolism, splicing, mitochondrial metabolism and lipid metabolism. Our study shows that RBM3 in skeletal muscle cells may define an important signaling hub that can be targeted both to improve metabolism and rejuvenate aged myoblasts.

Here we have shown that RBM3 directly controls the metabolism of muscle stem cells and their regenerative capacity *in vitro*. In the future, it will be important to determine the effect of RBM3 overexpressing myoblasts during skeletal muscle regeneration *in vivo*. The direct targets of RBM3 (both mRNA and protein) that mediate its effects remain less understood. It is possible that RBM3 (like other RNA binding proteins containing the RRM and disordered domains) may form aggregates like stress granules and it might control the lifetime and translation of specific RNAs involved in metabolism. The molecular mechanisms governing the choice and regulation of RNA targets of RBM3 are yet to be understood^41, 42^. Overall, controlling the expression of RBM3 using physiological conditions or small molecules presents an important avenue for regulating the stem cell state and metabolism.

## Materials and Methods

### Cell culture techniques

C2C12 cells (purchased from ATCC) were grown and maintained in DMEM (Dulbecco’s Modified Eagle Medium, Gibco-11995065) with 10% FBS (Foetal Bovine Serum, Gibco-16000044) and 1% Pen-Strep (Penicillin-Streptamycin, Gibco-15140122). Mouse primary myoblasts were cultured on ECM Gel coated dishes (Sigma-Aldrich-E6909). The cells were grown in 37^0^C and 5% CO_2_ conditions. C2C12 differentiation was done using DMEM + 2% horse serum (Gibco-26050070) and 1% Pen-Strep. Differentiation media was added once the cells reach 100% confluency and differentiation was done for 6 days. HEK293T cells were grown and maintained in DMEM with 10% FBS and 1% Pen-Strep. For hypothermic stress, cells were grown in 25^0^C and 32^0^C and 5% CO_2_ conditions for 72 hrs.

### Preparation of retrovirus and stable cell lines

For retrovirus production, HEK293T was cotransfected with the plasmid containing the gene of interest pMIG-GFP, pMIG-RBM3 and packaging plasmid pCL-ECO using jetPRIME reagent (Polyplus-101000046). Media was collected 48 hrs. and 72 hrs. after transfection. The collected media was centrifuged at 4000 RPM for 5 minutes at room temperature. The supernatant was collected and concentrated using Retro-X-Concentrator from Takara Biosciences (Takara #631455) overnight at 4^0^C. The following day the concentrate was centrifuged at 1500 RPM for 45 minutes at 4^0^C to obtain the virus pellet. The pellet was resuspended in DMEM media and aliquoted and stored at −80^0^C if not used immediately.

The retroviral titre calculation was done using the Retroviral titration kit from Takara Biosciences (Retro-X™ qRT-PCR Titration Kit 631453) using the qRT-PCR-based method. C2C12 cells were transduced with 10^5^-10^6^ virus particles. C2C12 cells were seeded in a 6 cm dish and 10^5^-10^6^ virus particles were added dropwise to media containing 8 ug/mL Sequabrene (Sigma S2667-1VL) and 25 mM Hepes buffer. The media was changed 24 hrs. after transduction. Cells were checked 48 hrs. after transduction and an efficiency of 90-95% was observed.

### si-RNA treatment on C2C12 cells

C2C12 cells were transfected with 200 nM of siRBM3 (Dharmacon L-041823-01-0005) and 200 nM scrambled control (Dharmacon D-001810-10-05) using dharmafect transfection reagent following dharmefect transfection protocol.

### Protein isolation and Western blotting

Protein isolation was done using RIPA buffer (Thermo Scientific-89901) and 1X PIC (Protease inhibitor cocktail cOmplete Roche 11697498001). The concentration of the protein was done using Bradford assay (Puregene PG-035-500mL) or BCA assay (G-Biosciences-786570). For western blotting 25 ug protein was loaded in each well. 3% BSA in TBST was used as a blocking buffer. Primary antibodies MyHC (Invitrogen 14650382), MF-20 (DHSB AB_2147781) MYOG (Invitrogen MA5-11486), MyoD1 (Santa Cruz SC-377460), RBM3 (Invitrogen PA5-51976), beta-ACTIN (CST 4967S), beta-Tubulin (CST 2146), PKM1(CST D30G6), beta-Actin (CST 4967) PKM2 (CST D78A4), PDH (CST 3205), SDHA (CST 5839), phospho-4E-BP1 (CST 2855), total 4E-BP1 (CST 9452), phospho-AMPK-alpha (CST 2535), total-AMPK-alpha (CST 5831), phospho-AMPK-beta (CST 4186), total-AMPK-beta (CST 4150), phospho-ACC (CST 11818) and total ACC (CST 3676) were used in 1:1000 dilution and incubated at 4^0^C overnight. Secondary antibodies: anti-mouse IgG, HRP-linked (CST 7076) and anti-rabbit IgG, HRP-linked (CST 7074) were used in 1:5000 dilution and incubated at room temperature for 1hr. The blots were developed using a developing solution (advansta-K-12049- D50).

### RNA isolation and qRT-PCR

RNA isolation from cell samples was done using Trizol (Thermo Scientific-15596018). Reverse transcription was done using the TAKARA cDNA synthesis kit (PrimeScript™ 1^st^ strand cDNA Synthesis Kit 6110A). qRT-PCR was done using SYBR-Green (PowerUp™ SYBR™ Green Master Mix A25742). Following primers were used for qRT:

*MyHC* (Forward primer: TAAACGCAAGTGCCATTCCTG, Reverse primer: GGGTCCGGGTAATAAGCTGG), *Myog* (Forward primer: CGATCTCCGCTACAGAGGC, Reverse primer: GTTGGGACCGAACTCCAGT), *MyoD1* (Forward primer: TCCGCTACATCGAAGGTCTG, Reverse primer: GTCCAGGTGCGTAGAAGGC), rbm3 (Forward primer: CTTCGTAGGAGGGCTCAACTT, Reverse primer: CTCCCGGTCCTTGACAACAAC), 18S rRNA (Forward primer: CCCGTTGAACCCCATTCGTG, Reverse primer: GGGCCTCACTAAACCATCCA), pkm1 (Forward primer: GCCGCCTGGACATTGACTC, Reverse primer: CCATGAGAGAAATTCAGCCGAG), *Pkm2* (Forward primer: CGCCTGGACATTGACTCTG, Reverse primer: GAAATTCAGCCGAGCCACATT), *Sdha* (Forward primer: GGAACACTCCAAAAACAGACCT, Reverse primer: CCACCACTGGGTATTGAGTAGAA), *Sdhc* (Forward primer: GCTGCGTTCTTGCTGAGACA, Reverse primer: ATCTCCTCCTTAGCTGTGGTT), *Pdhb* (Forward primer: AGGAGGGAATTGAATGTGAGGT, Reverse primer: ACTGGCTTCTATGGCTTCGAT), *Pax7* (Forward primer: CTCAGTGAGTTCGATTAGCCG, Reverse primer: AGACGGTTCCCTTTGTCGC), *Myf5* (Forward primer: CACCACCAACCCTAACCAGAG, Reverse primer: AGGCTGTAATAGTTCTCCACCTG)

### MTT assay

For the MTT assay, 5 mg/mL MTT reagent was used (Sigma M2128-500MG). Cells were seeded in a 96-well plate. For drug assay, 10000 cells were seeded per well. The following day, cells were treated with drugs oligomycin (Sigma 75351) and 2DG (Sigma D8375) at different concentrations: 50 nM, 100 nM, and 200 nM for oligomycin and 0.25 mM, 0.5 mM, 1 mM for 2DG for 48 hrs. After which cells were incubated with MTT reagent for 3 hrs. at 37^0^C in dark. Absorbance reading was taken at 570 nm using a plate reader. For the cell viability experiment, C2C12 stable lines: GFP, RBM3 have been seeded in 96 well plates 5,000 cells per well. MTT assay was done 48 hrs. after seeding and absorbance was measured at 570 nm.

### Seahorse Assay

Seahorse assay was done to measure the oxygen consumption rate (OCR) of the cells using Seahorse XFe24 bioanalyser machine. For that, the 24 well cell culture plates (Agilent Technologies 102340-100) were seeded with 18000 cells per well. DMEM XF base media (Agilent 102353-100) containing 1 mM sodium pyruvate (Sigma S8636-100ML), 2 mM glutamine (Sigma G7513-100ML) and 20 mM glucose (Qualigens Q15405) for measuring OCR was used. Cells were incubated in a non-CO_2_ incubator at 37^0^C for 45 minutes. OCR were measured in response to the following drugs: 1 uM oligomycin, 3 uM FCCP (Cayman chemical 800364-9897), 1.5 uM of antimycin (Cayman chemicals 1397-94-0) and rotenone (Sigma R8875-1G) for OCR. Apart from the drug treatment, OCR was measured at the baseline levels.

### Proteomics sample extraction and preparation

C2C12 cells were subjected to hypothermic conditions: 32^0^C and 5% CO_2_ for different time points of 6 hrs., 12 hrs., 24 hrs., 48 hrs. C2C12 stable cell lines: GFP and RBM3 were used for proteomics studies. Cells were lysed with RIPA + Protease inhibitor cocktail. They were centrifuged at 10000 RPM for 30 minutes at 4^0^C. The supernatant was collected, and a BCA assay was done. 100 ug of protein was used for proteomic analysis. Chilled acetone was added to the protein samples and incubated for 3 hrs. at −80^0^C. Centrifugation was done at 13,000 RPM for 30 minutes at 4^0^C. The pellet was washed with acetone at 13000 RPM for 10 minutes at 4^0^C. The samples were treated with 8 M urea (in Ammonium bicarbonate) and 0.5 M DTT and incubated for 30 minutes at 37^0^C. Following this, the samples were treated with 30 mM iodoacetamide and incubated for 30 minutes at 37^0^C. The samples were then digested with trypsin (1:20 trypsin: protein) at 37^0^C overnight. The digested samples were desalted using the C18 column. The column was equilibrated with 0.1% formic acid. The sample was eluted in 100% Acetonitrile containing 0.1% formic acid. The eluted samples were dried using a speed vacuum.

### Separation and digested protein for Identification and quantitation of the Proteome using Sciex 7600 Zeno-TOF LCMS

Equal concentrations of protein samples were digested using mass spec grade trypsin and digested peptides were desalted using a C18 SPE cartridge. Peptides were analyzed using a Sceix Zeno TOF 7600 (AB Sciex, Foster City, CA, United States) equipped with a Shimadzu LC 40 UPLC system. Peptides were separated on Aquity UPLC BEH C18 column (150 × 1 mm, 1.7 µm,) column using a gradient of water and acetonitrile. For protein quantitation, spectral libraries were generated using information-dependent acquisition (IDA) mode after injecting 3 µg of tryptic digest on the above-mentioned C18 column using a Shimadzu LC40 autosampler system coupled with Sciex Zeno Tof 7600 fitted. For library generation, the peptides from all the experiment sets were pooled and injected into the mass spec for comprehensive coverage. The total chromatography run time was 180 minutes. The first 10 minutes of the column flow was sent to waste after loading for online desalting of the sample. A 180 minutes gradient in multiple steps (ranging from 5 to 30% acetonitrile in water containing 0.1% formic acid) was set up to elute the peptides from the column for the first 135 minutes then in the next 10 minutes concentration of B was increased to 55% and most of the peptide eluted till this time of 145 minutes. The concentration of mobile phase B was increased to 95% B in 6 minutes and held at this concentration for 5 minutes then the column was again kept at 98% of mobile phase A (Water and 0.1% formic acid) for 24 minutes for equilibration. The flow rate of the UPLC was set at 30µL/minutes. Three biological replicates of the samples were run for each experimental set. For library preparation the mass spec was run in an information-dependent acquisition mode (IDA) with the MS1 mass range of 350 Da to 1500 Da, Gas 1 was 25 L/min, Gas 2 was set at 15 L/minutes, whereas curtain gas was set at 25 L/minutes, ionization voltage was 5500 volt, accumulation time was set at 0.2 seconds and source temperature was at 350^0^C. The most abundant top 50 multiple charge precursor was fragmented in each cycle with dynamic collision energy in collision-induced dissociation (CID). The mass range for MS/MS was set at 150 to 1800 Da. Zeno trapping was on for the IDA data.

### Sequential Window Acquisition of all Theoretical Fragment Ion Spectra (SWATH) Analysis for Label-Free Quantification

For label-free quantification (SWATH analysis) the Q1 transmission windows were set to 25 Da from the mass range of 350 Da to 1500 Da. A total of 79 windows were acquired independently with an accumulation time of 50 milliseconds. The total cycle time was kept constant at 2 seconds. Protein Pilot^TM^ v. 5.0.2 was used to generate the spectral library. For label-free quantification, peak extraction and spectral alignment were performed using PeakView^R^ 2.2.0.11391 Software with the parameters set as follows: number of peptides, 2; number of transitions, 5; peptide confidence, 95%; XIC width, 20 ppm; XIC extraction window, 5 minutes. The data were further processed in MarkerView software ^TM^. 1.3.1 (AB Sciex, Foster City, CA, United States) for statistical data interpretation. In MarkerView^TM^, the peak areas under the curve (AUC) for the selected transition were normalized using total area sum intensities and all the biological replicates were averaged out before normalization. t-test was performed on the data set and fold-changed values were calculated for all the proteins.

### Data Analysis for proteomics

Data were processed with Protein Pilot Software v. 5.0.2 (ABSciex, Foster City, CA, United States) utilizing the Paragon and Progroup Algorithm. The analysis was done using the tools integrated into Protein Pilot at a 1% false discovery rate (FDR) with statistical significance. In brief, the UniProt mouse proteome database (UP000000589) was used to search for the matched peptide for library generation. The download included total combined (reviewed and un-reviewed) entries of 55,286 proteins. The resultant search identification file was used as a library for the extraction of peptide quantitation information from the SWATH acquisition. The extracted peptide information was processed using Marker view software ^TM^(V1.3) for statistical analysis. Biological triplicate data for each sample were normalized using the total area sum method and all the biological replicates were averaged out for the calculation of fold change calculation. After normalization t-test and principal component analysis were performed on the data set to check the possible correlated variables within the group. A volcano plot was generated to calculate the statistically significant fold change vs p-value. Proteins with less than 0.8-fold change were considered as downregulated and proteins with more than 1.5 fold change were considered as upregulated in the experiment sets. In this analysis library was made for approximately 2394 proteins that were identified with 0.05% False Discovery Rate (FDR) and approximately 1400 proteins were quantified in the data set. GO enrichment analysis, process and pathway enrichment analysis were done using metascape software.

### Metabolite Extraction and mass spectrometry

C2C12 stable lines-GFP, RBM3 were seeded in 10 cm cell culture dishes. Metabolite extraction was done using chilled 50% methanol^43^. The cells were scrapped in 50% methanol and 100 ug D4 alanine (Cambridge isotope lab DLM-250-1) and centrifuged at 13000 RPM for 5 minutes at room temperature. The supernatant was dried using a speed vacuum. The dried sample was stored at – 80^0^C. The pellet was resuspended in RIPA and protein was prepared and a BCA assay was done to determine the concentration. The media from the cells were processed similarly using 50% methanol. The dried samples were reconstituted and run on AB Sciex 5500 using the synergi rp fusion column.

For the detection of glycolytic and pentose phosphate pathway intermediates along with NAD, NADH, and acetyl-CoA, the samples were analysed in a triple-quadrupole type mass spectrometer (Sciex QTRAP 5500) with Schmadzu HPLC unit. Multiple reaction monitoring (MRM) methods were modified as published by *Walvekar et al* (2018) The metabolites were measured using Synergi, 4 μm Fusion-RP 80 A (150 x 2 mm; Phenomenex # 00F-4424-B0) by the methods described below: Samples were reconstituted in 100 μl of 50% methanol. Synergi, 4 μm Fusion-RP 80 A (150 x 2 mm; Phenomenex # 00F-4424-B0) Pump B; 0 minute: 0%; 2.0 minutes: 5%; 6.7 minutes: 60%; 7.3 minutes: 95%; 9.3 minutes: 95%; 10 minutes: 5%; 10.3 minutes: 0%; 12 minutes: 0%; 12.1 minutes: controller stop; flow rate 0.2 ml/minutes; autosampler temperature 5^0^C; Mobile phase A: 5mM Ammonium Acetate (Sigma 73594-25G-F) pH 8.0; Mobile phase B:100% acetonitrile (Fischer Scientific A955-4)

The area under each peak was calculated using AB SCIEX MultiQuant software 3.0.1

For TCA intermediates along with lactate and pyruvate, the mobile phase consisted of a premixed ratio of Water/Methanol (95:5) with 0.2 % Formic acid. The column heating oven was set to 45^0^C, autosampler temperature 5^0^C; flow rate 0.4 mL/minutes. The MS method was used as such as per the manufacturer’s instruction (Phenomenex TN-1241).

### Lipidomics sample preparation and analysis

C2C12 stable lines-GFP, RBM3 were seeded in a 6 cm dish. Lipid extraction was done using the SIMPLEX protocol ^44^. 225 uL of methanol containing 10 uL SPLASH LIPIDOMIX standard (Avanti polar lipids 330707) was added to the cell pellet and transferred to lo-bind Eppendorf tubes. The tubes were vortexed for 20 seconds and incubated in liquid nitrogen for 1 minute. It was then thawed at room temperature and sonicated for 30 minutes. 750 uL of MTBE (Sigma 650560) was added to the vials and vortexed for 1 hr. at 4^0^C. 188 uL of water was added to induce phase separation after which centrifugation was done at 10,000g for 5 minutes at 4^0^C. The upper phase (600 uL) was collected for lipidomics. The solvent was dried and resuspended in 200 uL of solvent B (10 mM ammonium formate, 0.1% formic acid in 90:10, 2-Propanol: acetonitrile). For the liquid chromatography separation, mobile phase A was water/acetonitrile (60/40, v/v) and mobile phase B was 2-propanol/acetonitrile (90/20, v/v), with both containing 10 mM ammonium formate and 0.1% formic acid. A 5 uL sample, thermostatted to 10^0^C, was injected onto a Thermo Scientific, Acclaim C30, reversed phase column (3 µm, 2.1 × 100 mm) thermostatted to 40^0^C in a Thermo Vanquish UHPLC system. Gradient elution was performed at a flow rate of 300 µL/minutes, beginning with 20% mobile phase B, that was increased to 40% B over 1 minutes, to 80% B over 8 minutes, to 100% B over 11 minutes, and held at 100% B until 12 minutes, the column was re-equilibrated at 20% mobile phase B from 13 minutes to 15 minutes, giving a total run time of 15 minutes. The lipidomic analysis was performed on a quadrupole-orbitrap mass spectrometer (Q Exactive, Thermo Fisher Scientific) operating in positive ion mode via electrospray ionization and used to scan from m/z 150 to 1000 at 1 Hz and 140,000 resolution. Data were analyzed using the Lipid Search software.

### Immunofluorescence and confocal microscopy

C2C12 cells were seeded on coverslips and subjected to hypothermia (32^0^C) and control (37^0^C) for 72 hrs. Cells were fixed with 4% PFA for 15 minutes at room temperature. Permeabilization was done using 0.1% triton-X-100 in 1X-PBS for 10 minutes at room temperature. Blocking was done for 1 hr. using 5% BSA in PBST at room temperature. Primary antibody (RBM3, Ki-67 (Invitrogen PA5-19462)) was used at a dilution of 1:300 and incubated at 4^0^C overnight. The secondary antibody Alexa fluor 647 goat anti-rabbit (Invitrogen A21245) was used at 1:500 dilution for 1 hr. at room temperature. For nuclear staining, DAPI (Thermo Scientific 62248) was used at a dilution of 1:1000 for 5 minutes. Coverslips were mounted using Prolong gold mounting agent (Life technologies P36930). Images were taken using a confocal microscope.

### Mitotracker and TMRM staining

C2C12 stable lines-GFP, RBM3 were seeded in imaging dishes. After 24 hrs., the cells were stained with mitotracker (Invitrogen M7512) for 15 minutes at 37^0^C. The cells were washed with PBS. For nuclear staining, cells were incubated with Hoechst (Invitrogen H3570) (1:1000) for 5 minutes and cells were imaged using a confocal microscope. For TMRM staining, cells were incubated with TMRM (Sigma T5428-25MG) stain for 10 minutes at 37^0^C and Hoechst for 5 minutes and imaged using a confocal microscope.

### Neon transfection

Mouse primary myoblasts were transfected with pMIG-RBM3 and pMIG-GFP using the Neon TM transfection system (Invitrogen MPK 1025). The cells were trypsinized and counted using a hemocytometer. For 12 well plate 3,00,000 cells and 2 ug DNA was used for transfection. The cell pellet was resuspended in 10 uL Buffer R and DNA was added to it. The cell suspension with DNA was placed on the neon tube containing 3mL of Buffer E and the electroporation was performed using the transfection program. The parameters used for transfection are voltage: 1500V, pulse width: 10 milliseconds, no of pulse:3. The cells were grown without antibiotics for 48 hrs. after transfection. After 48 hrs. cells were grown in growth media containing antibiotics.

### Satellite cell isolation and growth techniques

Satellite cells were isolated from young (3-4 months) and aged (18 months) BL6/J mice using the Miltenyi satellite cell isolation kit (Miltenyi biotech 130-104-268). Due to the low viability of cells using the kit-based method, a non-kit-based method was used to isolate satellite cells from aged mice^45^. Before seeding, the tissue culture dishes were coated with ECM.

### Cloning techniques

Mouse RBM3 gene from RBM3-PUC plasmid (Origene MC203679) was cloned into pMIG-GFP plasmid by restriction digestion method using Bgl2 and EcoR1 (NEB). Primers used for cloning (Forward primer: GGAAGATCTTATGTCGTCTGAAGAAGGGAAACTC, Reverse primer: CCGGAATTCTCAGTTGTCATAATTGTCTCT). ΔRBD was created by deleting 143 bp from the N terminus of the RBM3 sequence. ΔRBD was cloned into pMIG-GFP plasmid by restriction digestion method using Bgl2 and EcoR1 using the following primers: Forward primer: GGAAGATCtATGTTTGGCTTCATCACC, Reverse primer: CCGGAATTCTCAGTTGTCATAATTGTC. The clone sequences were verified using the Sanger sequencing technique. All the plasmids were purified using Qiagen miniprep kit (Qiagen 27104). For neon transfection of primary myoblasts, RBM3-GFP plasmid from Origene was used (Origene MG201130).

### Statistical Analysis

All the data are represented as mean ± SEM. All the bar graphs were generated using graph pad Prism 8 using unpaired t-test or two-way ANOVA using multiple comparisons. Statistical significance was accepted at p <= 0.05

## Acknowledgements

The authors would like to acknowledge funding from the Department of Biotechnology, Govt of India. PD was funded by the CSIR fellowship, SM was funded by the DBT-RA fellowship. We thank Prof. Apurva Sarin for pMIG-GFP plasmid.

## Author Contributions

PD contributed to the cell culture, molecular biology, biochemical assays and writing of the manuscript. SR and PST contributed to the cell culture and molecular biology work. NS and PS contributed to western blot analysis. MAH performed metabolomic analysis. SM, RGHM performed Lipidomic analysis, HL and SSP performed confocal imaging. NS contributed to the proteomic studies. AR participated in the conceptualization and writing of the manuscript.

## Data sharing plans

All raw data including mass spectrometry data will be shared in publicly available databases upon publication. All the mass spec raw and processed data file will be uploaded on the Proteomics identification database (PRIDE) for review and accessibility. Other data file of the experiment will also be available as made available for review.

## Additional information

Supplementary information is available for this paper †All correspondence addressed to Arvind Ramanathan **Email:**

## Supplementary figure legends

**Supplementary 1 (A)** mRNA expression levels of Myosin heavy chain (*MyHC*) during differentiation using C2C12 cells at 25^0^C and 32^0^C compared to each other after 72 hrs. of hypothermic adaptation where the x-axis represents the number of days pre-differentiation and during differentiation and the y-axis represents the mRNA fold change of *MyHC* **(B)** Representative images of C2C12 myoblasts taken under bright field microscope at 10X. (Scale bar, 50 um) and myotubes (Scale bar, 100 um).

**Supplementary 2 (A)** Heat map of upregulated proteins involved in mitochondrial and fatty acid metabolism at different time points of 6 hrs., 12 hrs., 24 hrs. and 48 hrs. (n=3). **(B)** Heat map of upregulated proteins involved in RNA processing and metabolism at different time points of 6 hrs., 12 hrs., 24 hrs. and 48 hrs. (n=3). **(C)** Volcano plot of 37^0^C compared to 32^0^C hypothermia at 6 hrs., 12 hrs., 24 hrs., 48 hrs., RBM3 overexpressed and GFP control where x-axis represents p-value and y-axis represents log (fold change).

**Supplementary 3 (A)** Bar graph representing the total number of upregulated and downregulated proteins and % of upregulated and downregulated proteins at different time points of 6 hrs., 12 hrs., 24 hrs. and 48 hrs. at 32^0^C. GO enrichment pathway analysis of upregulated proteins at **(B)** 6 hrs. **(C)** 12 hrs. **(D)** 24 hrs. and **(E)** 48 hrs. at 32^0^C as analyzed by metascape.

**Supplementary 4 (A)** Western blot analysis of RBM3 using different concentrations of siRBM3 in C2C12 cells at 37^0^C for 72 hrs. **(B)** mRNA expression levels of RBM3 using C2C12 cells overexpressing pMIG-GFP control, pMIG-RBM3 at 37^0^C (n=2). **(C)** mRNA expression levels of *MyoD1* in mouse primary myoblasts transfected with pMIG-RBM3 and pMIG-GFP (n=2). **(D)** Representative images of stable C2C12 cells transduced with pMIG-RBM3 in bright field and epifluorescence (Scale bar, 100 um) **(E)** Epifluorescence images of C2C12 transfected with RBM3-GFP taken at 100X (oil) (Scale bar, 16 um). *, **, *** represents p-value < 0.05, 0.01 and 0.001 respectively.

**Supplementary 5 (A)** GO enrichment analysis of protein functional network of proteins upregulated in C2C12 cells overexpressing RBM3 analyzed by Metascape.

**Supplementary 6 (A)** Bar graph showing the % viability of C2C12 cells overexpressing pMIG-GFP, pMIG-RBM3 in presence of different concentrations of oligomycin (50 nM, 100 nM and 200 nM) **(B)** IC50 plot for oligomycin using C2C12 cells overexpressing pMIG-GFP control and pMIG-RBM3 where the x-axis represents the oligomycin concentration and the y-axis represents the % viability of the cells (n=2). **(C)** Bar graph showing the % viability of C2C12 cells overexpressing pMIG-GFP control, pMIG-RBM3 in presence of different concentrations of 2DG. **(D)** IC50 plot for 2DG using C2C12 cells overexpressing pMIG-GFP control and pMIG-RBM3 where the x-axis represents the oligomycin concentration and the y-axis represents the % viability of the cells (n=2). *, **, *** represents p-value < 0.05, 0.01 and 0.001 respectively.

**Supplementary 7 (A)** Confocal images of C2C12 cells overexpressing pMIG-GFP control and pMIG-RBM3. Red indicates mitotracker (100 mM), blue indicates Hoescht and green indicates GFP (Scale bar, 16 um) **(B)** Confocal images of C2C12 cells overexpressing pMIG-GFP control and pMIG-RBM3. Red indicates TMRM, blue indicates Hoescht and green indicates GFP (Scale bar, 16 um) **(C)** Western blot analysis of SDHA and PDH using C2C12 cells overexpressing pMIG-GFP control and pMIG-RBM3 **(D)** mRNA expression levels of *Sdha*, *Sdhc* and *Pdhb* using C2C12 cells overexpressing pMIG-GFP control and pMIG-RBM3 (n=4). **(E)** mRNA expression levels of *Sdha*, *Sdhc* and *Pdhb* using mouse primary myoblasts overexpressing pMIG-GFP control and pMIG-RBM3 (n=3). **(F)** Heat map showing levels of lactate/pyruvate using C2C12 cells overexpressing pMIG-GFP control and pMIG-RBM3 (n=3). **(G)** Heat map showing levels of lactate/pyruvate using media from C2C12 cells (48 hrs.) overexpressing pMIG-GFP control and pMIG-RBM3 (n=3). **(H)** Western blot analysis of AMPK-alpha using C2C12 cells overexpressing pMIG-GFP control and pMIG-RBM3 **(I)** Bar graph quantifying the levels of phosphorylated/total AMPK-alpha. *, **, *** represents p-value < 0.05, 0.01 and 0.001 respectively.

**Supplementary 8** Volcano plot of **(A)** Intracellular metabolites of C2C12 cells overexpressing GFP and RBM3 **(C)** Extracellular metabolites of C2C12 cells overexpressing GFP and RBM3 where the x-axis represents the log of fold change and the y-axis represents -log of p-value. Grey dots represent non-significant metabolite species and blue dot represents significantly high levels of metabolite species (n=3). Heat map showing levels of TCA metabolites using C2C12 cells overexpressing pMIG-GFP control and pMIG-RBM3 **(B)** intracellular **(C)** extracellular (n=3). *, **, *** represents p-value < 0.05, 0.01 and 0.001 respectively.

**Supplementary 9 (A)** Volcano plot of GFP and RBM3 overexpression where the x-axis represents the log of fold change and the y-axis represents -log of p-value. Grey dots represent non-significant lipid species and blue dot represents significantly low levels of lipid species (n=3). **(B)** Heat map showing levels of cholesterol esters and triglycerides using C2C12 cells overexpressing pMIG-GFP control and pMIG-RBM3 (n=3). *, **, *** represents p-value < 0.05, 0.01 and 0.001 respectively.

**Supplementary 10** mRNA expression levels of **(A)** *MyHC* and **(B)** *Myog* of aged and young myoblasts (n=3). **(C)** mRNA expression levels of *Myog* of aged myoblasts at 32^0^C compared to 37 ^ͦ^ C (n=2). **(D)** Bar graph showing the comparative ΔCt values of young primary, aged primary and aged primary transfected with RBM3-GFP (n=2). Heat map showing levels of extracellular lactate/pyruvate ratio **(E)** 24 hrs. **(F)** 48 hrs. and TCA metabolites in aged mouse primary myoblasts transfected with RBM3-GFP and GFP control **(G)** 24 hrs. **(H)** 48 hrs. (n=3). *, **, *** represents p-value < 0.05, 0.01 and 0.001 respectively.

**Supplementary 11 (A)** Representative images of mouse primary myoblasts (young and aged) and differentiated cells (Scale bar, 100 um) **(B)** Representative images of mouse primary young myoblasts transfected with pMIG-RBM3 and pMIG-GFP in bright field and epifluorescence (Scale bar, 100 um).

