## Supplementary file for "PROTEOMICS OF HYPOTHERMIC ADAPTATION REVEALS THAT RBM3 ENHANCES MITOCHONDRIAL METABOLISM AND MUSCLE STEM-CELL DIFFERENTIATION"

**Supplementary Fig 1**

S1 (A)

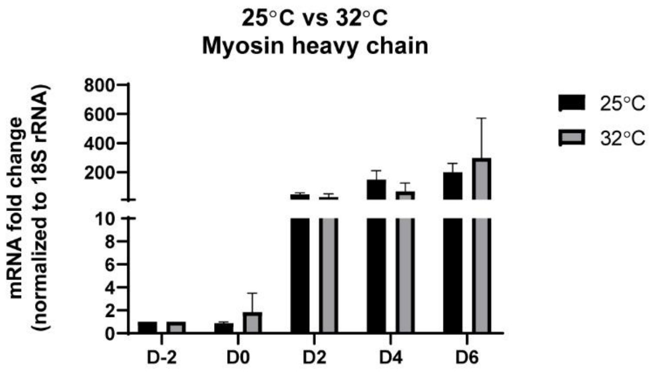


S1 (B) REPRESENTATIVE IMAGES OF C2C12 MYOBLAST AND DIFFERENTIATION

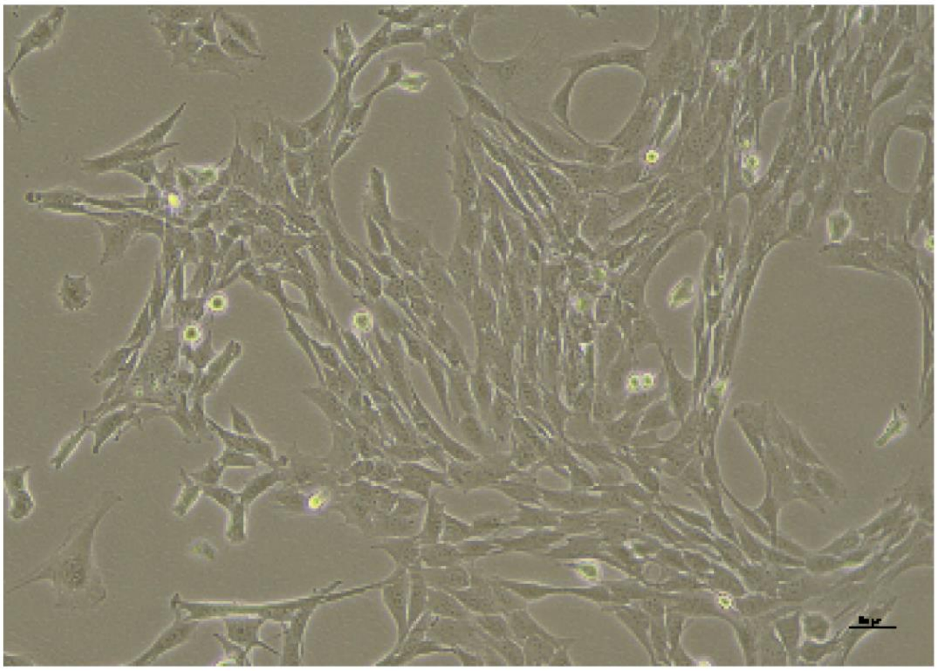

Myoblasts

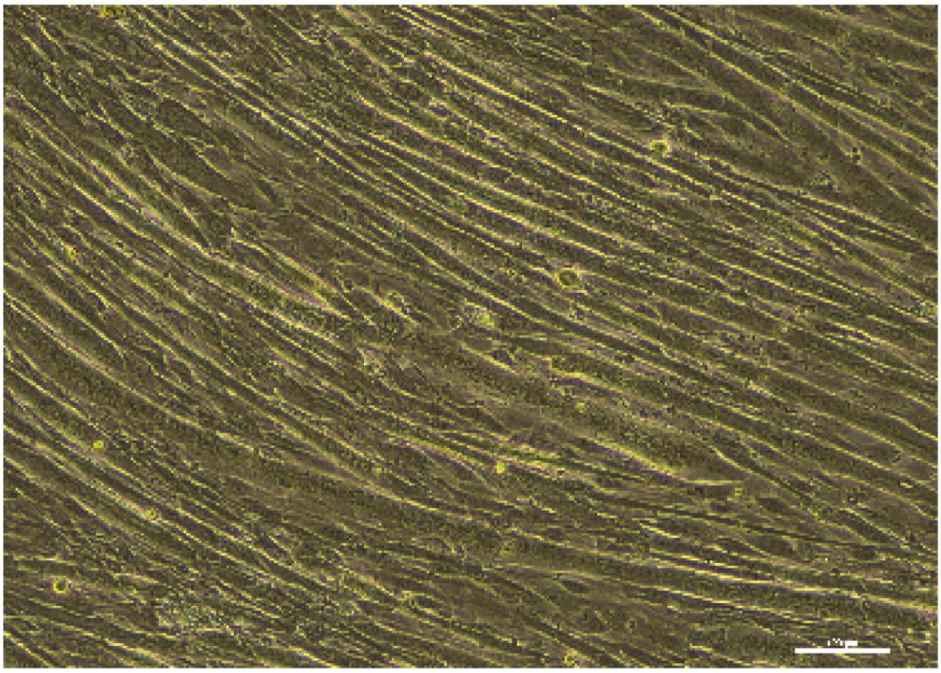

Myotubes

**Supplementary 1 (A)** mRNA expression levels of Myosin heavy chain (*MyHC*) during differentiation using C2C12 cells at $25^0$C and $32^0$C compared to each other after 72 hrs. of hypothermic adaptation where the x-axis represents the number of days pre-differentiation and during differentiation and the y-axis represents the mRNA fold change of *MyHC* **(B)** Representative images of C2C12 myoblasts taken under bright field microscope at 10X. (Scale bar, 50 um) and myotubes (Scale bar, 100 um).

Supplementary Fig 2

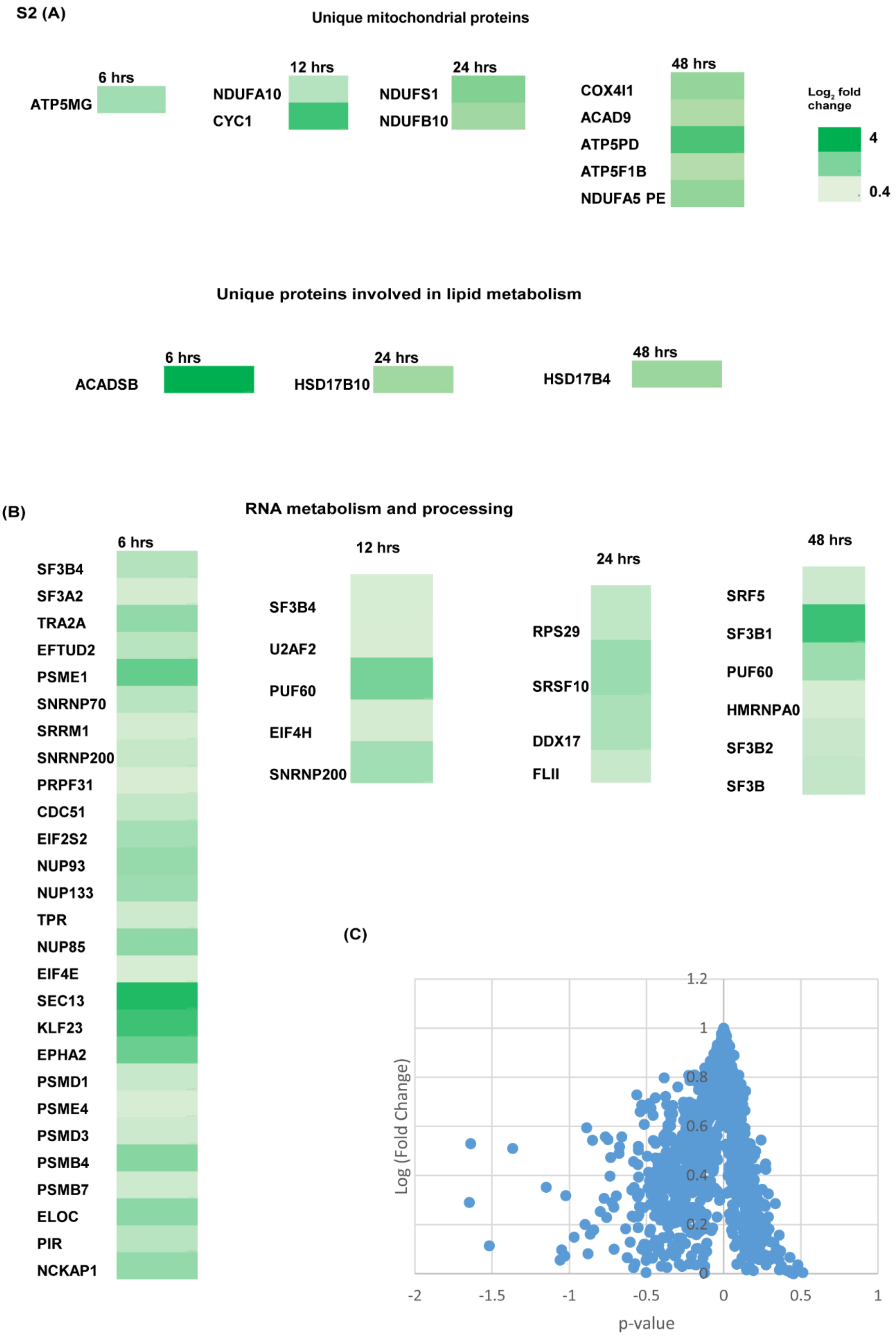


**Supplementary 2 (A)** Heat map of upregulated proteins involved in mitochondrial and fatty acid metabolism at different time points of 6 hrs., 12 hrs., 24 hrs. and 48 hrs. (n=3). **(B)** Heat map of upregulated proteins involved in RNA processing and metabolism at different time points of 6 hrs., 12 hrs., 24 hrs. and 48 hrs. (n=3). **(C)** Volcano plot of $37^0$C compared to $32^0$C hypothermia at 6 hrs., 12 hrs., 24 hrs., 48 hrs., RBM3 overexpressed and GFP control where x-axis represents p-value and y-axis represents log (fold change).

**Supplementary Fig 3**

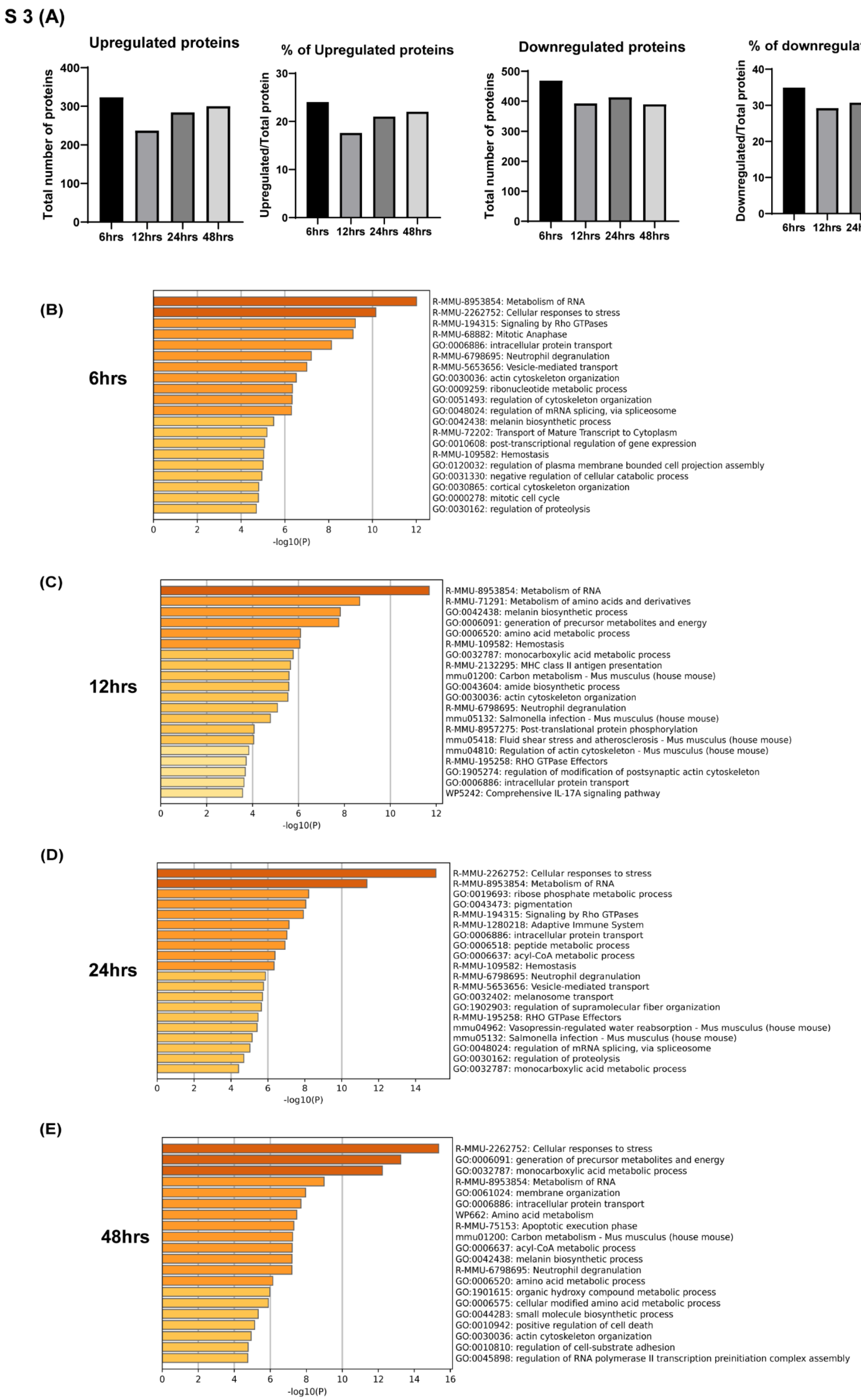

**Supplementary 3 (A)** Bar graph representing the total number of upregulated and downregulated proteins and % of upregulated and downregulated proteins at different time points of 6 hrs., 12 hrs., 24 hrs. and 48 hrs. at $32^0$C. GO enrichment pathway analysis of upregulated proteins at **(B)** 6 hrs. **(C)** 12 hrs. **(D)** 24 hrs. and **(E)** 48 hrs. at $32^0$C as analyzed by metascape.

**Supplementary Fig 4**

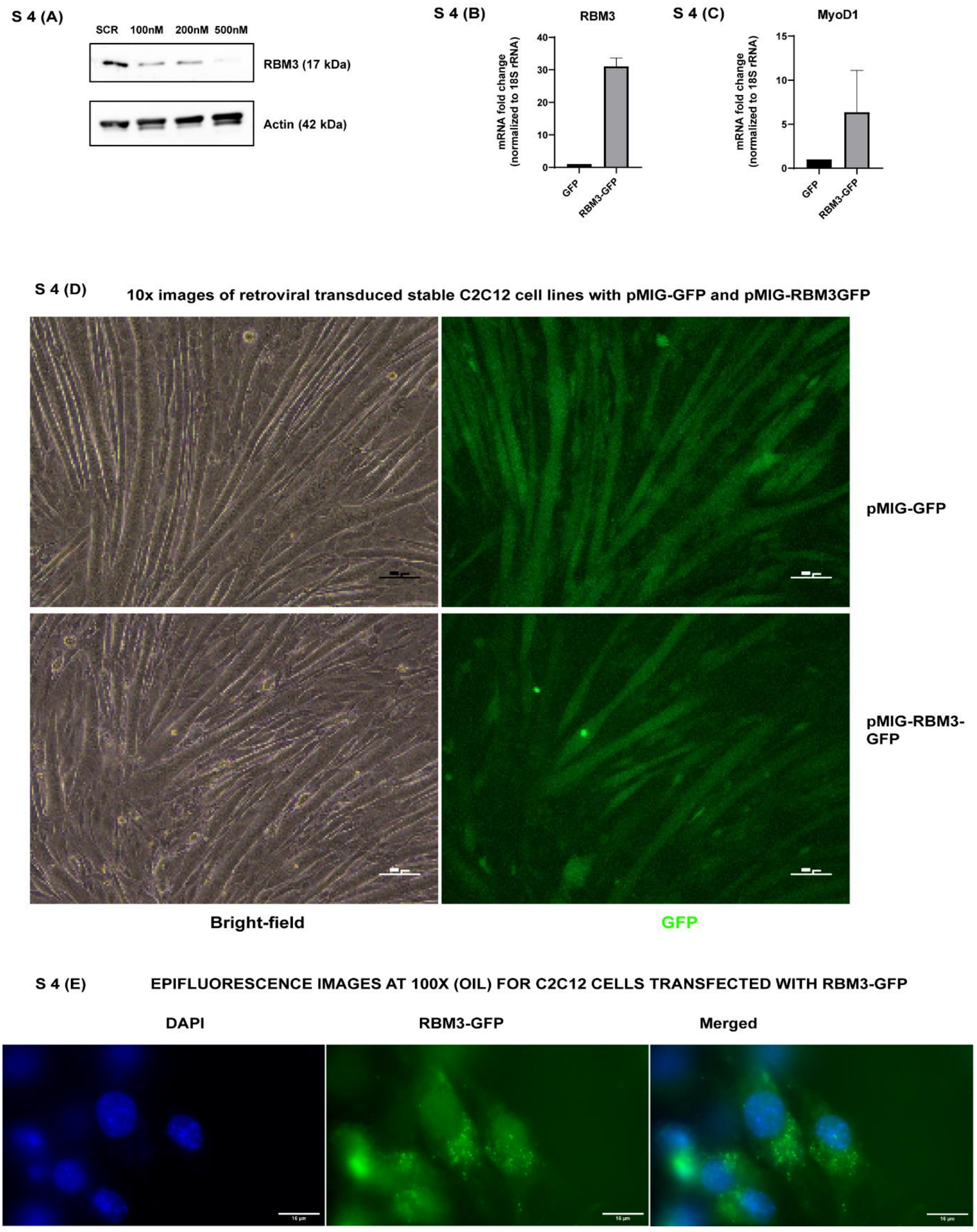


**Supplementary 4 (A)** Western blot analysis of RBM3 using different concentrations of siRBM3 in C2C12 cells at 37$^0$C for 72 hrs. **(B)** mRNA expression levels of RBM3 using C2C12 cells overexpressing pMIG-GFP control, pMIG-RBM3 at 37$^0$C (n=2). **(C)** mRNA expression levels of *MyoD1* in mouse primary myoblasts transfected with pMIG-RBM3 and pMIG-GFP (n=2). **(D)** Representative images of stable C2C12 cells transduced with pMIG-RBM3 in bright field and epifluorescence (Scale bar, 100 um) **(E)** Epifluorescence images of C2C12 transfected with RBM3-GFP taken at 100X (oil) (Scale bar, 16 um). *, **, *** represents p-value < 0.05, 0.01 and 0.001 respectively.

**Supplementary Fig 5**

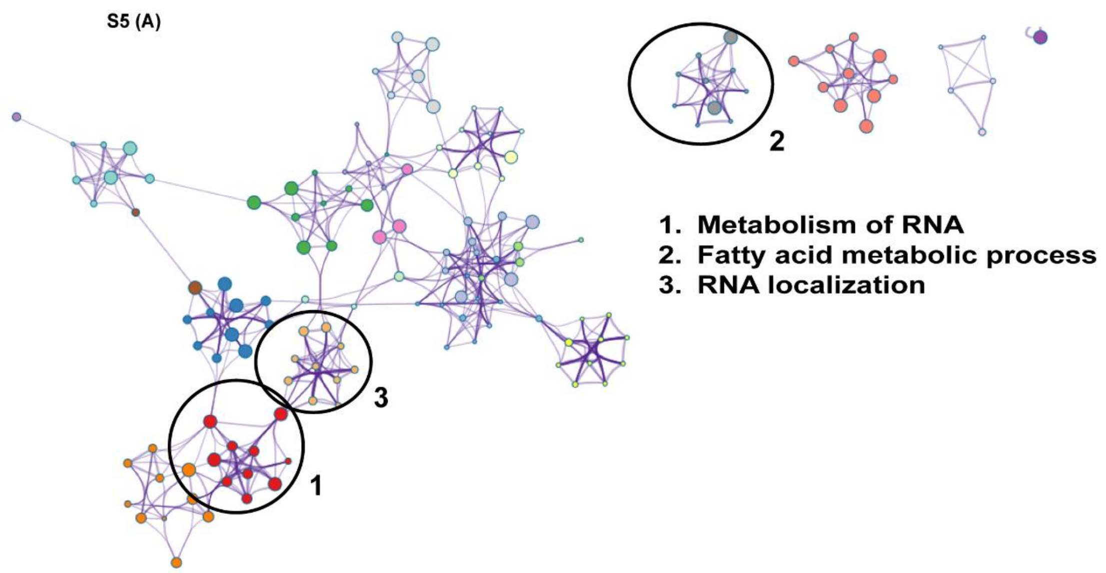


**Supplementary 5 (A)** GO enrichment analysis of protein functional network of proteins upregulated in C2C12 cells overexpressing RBM3 analyzed by Metascape.

**Supplementary Fig 6**

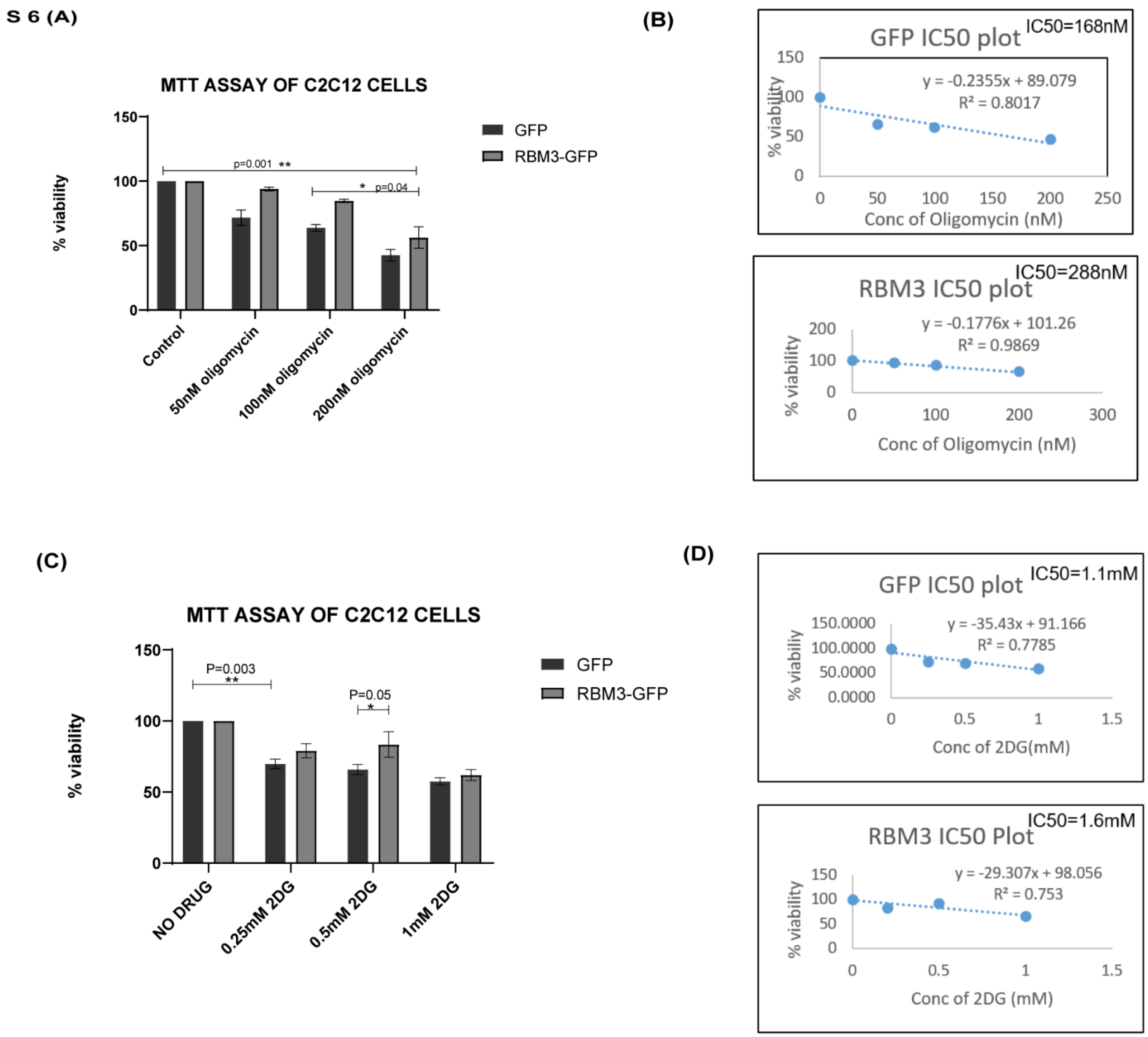


**Supplementary 6 (A)** Bar graph showing the % viability of C2C12 cells overexpressing pMIG-GFP, pMIG-RBM3 in presence of different concentrations of oligomycin (50 nM, 100 nM and 200 nM) **(B)** IC50 plot for oligomycin using C2C12 cells overexpressing pMIG-GFP control and pMIG-RBM3 where the x-axis represents the oligomycin concentration and the y-axis represents the % viability of the cells (n=2). **(C)** Bar graph showing the % viability of

C2C12 cells overexpressing pMIG-GFP control, pMIG-RBM3 in presence of different concentrations of 2DG. **(D)** IC50 plot for 2DG using C2C12 cells overexpressing pMIG-GFP control and pMIG-RBM3 where the x-axis represents the oligomycin concentration and the y-axis represents the % viability of the cells (n=2). *, **, *** represents p-value < 0.05, 0.01 and 0.001 respectively.

Supplementary Fig 7

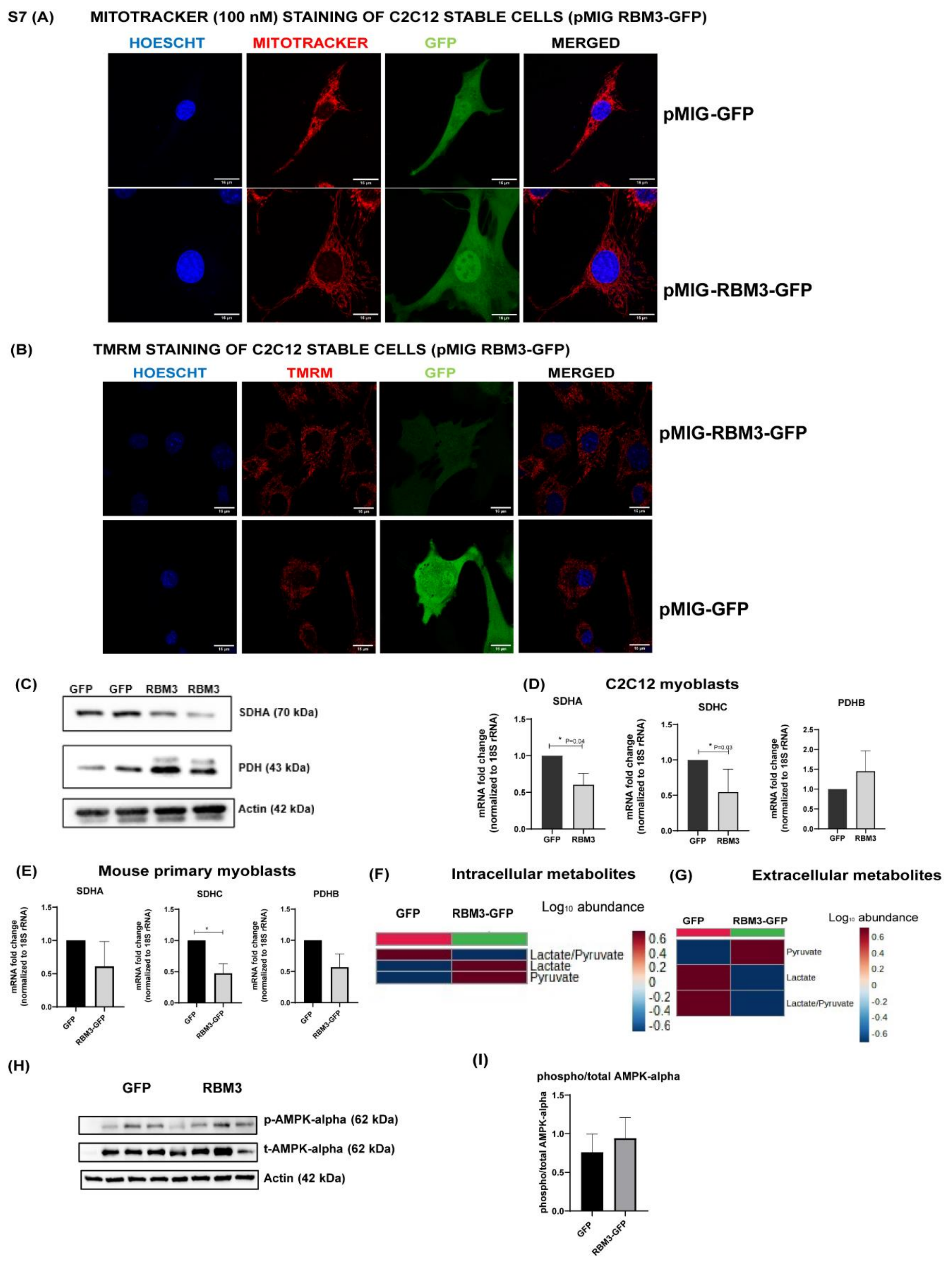


**Supplementary 7 (A)** Confocal images of C2C12 cells overexpressing pMIG-GFP control and pMIG-RBM3. Red indicates mitotracker (100 mM), blue indicates Hoescht and green indicates

GFP (Scale bar, 16 um) **(B)** Confocal images of C2C12 cells overexpressing pMIG-GFP control and pMIG-RBM3. Red indicates TMRM, blue indicates Hoescht and green indicates GFP (Scale bar, 16 um) **(C)** Western blot analysis of SDHA and PDH using C2C12 cells overexpressing pMIG-GFP control and pMIG-RBM3 **(D)** mRNA expression levels of *Sdha*, *Sdhc* and *Pdhb* using C2C12 cells overexpressing pMIG-GFP control and pMIG-RBM3 (n=4). **(E)** mRNA expression levels of *Sdha*, *Sdhc* and *Pdhb* using mouse primary myoblasts overexpressing pMIG-GFP control and pMIG-RBM3 (n=3). **(F)** Heat map showing levels of lactate/pyruvate using C2C12 cells overexpressing pMIG-GFP control and pMIG-RBM3 (n=3). **(G)** Heat map showing levels of lactate/pyruvate using media from C2C12 cells (48 hrs.) overexpressing pMIG-GFP control and pMIG-RBM3 (n=3). **(H)** Western blot analysis of AMPK-alpha using C2C12 cells overexpressing pMIG-GFP control and pMIG-RBM3 **(I)** Bar graph quantifying the levels of phosphorylated/total AMPK-alpha. *, **, *** represents p-value < 0.05, 0.01 and 0.001 respectively.

Supplementary Fig 8

S8 (A)

Intracellular metabolites

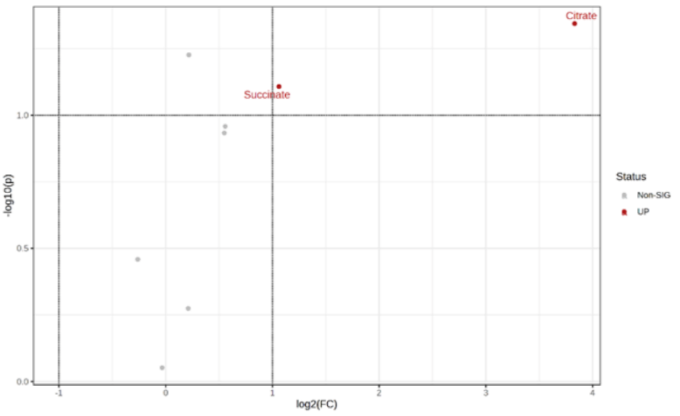


(B)

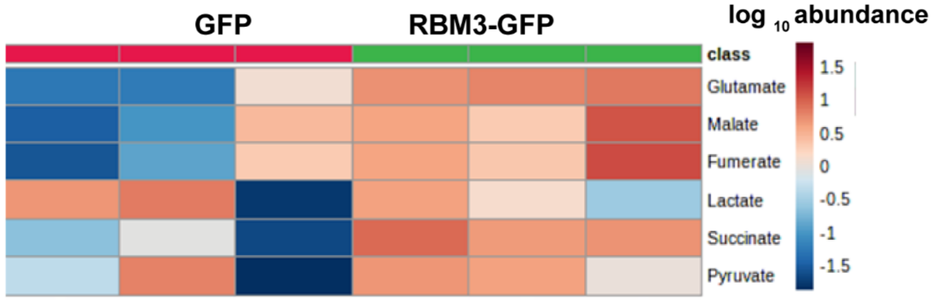


Extracellular metabolites

(C)

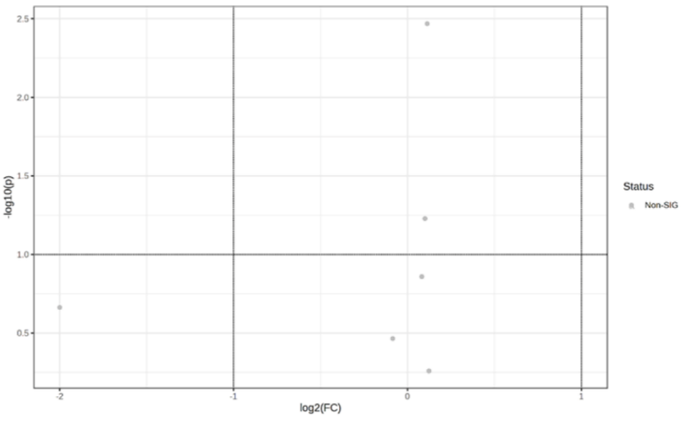


(D)

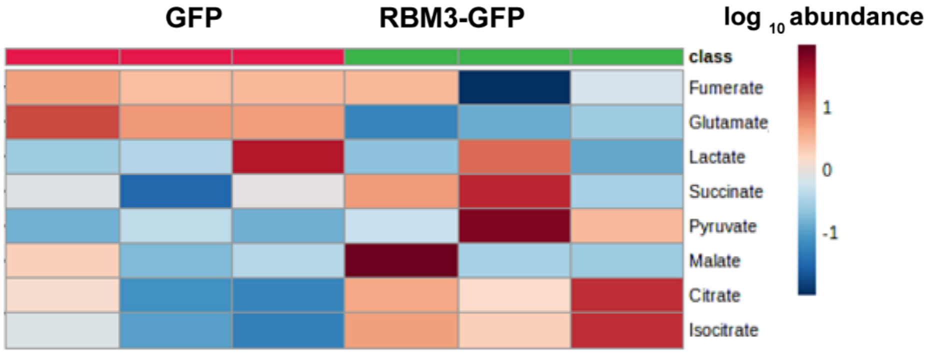

**Supplementary 8** Volcano plot of **(A)** Intracellular metabolites of C2C12 cells overexpressing GFP and RBM3 **(C)** Extracellular metabolites of C2C12 cells overexpressing GFP and RBM3 where the x-axis represents the log of fold change and the y-axis represents -log of p-value . Grey dots represent non-significant metabolite species and blue dot represents significantly high levels of metabolite species (n=3). Heat map showing levels of TCA metabolites using C2C12 cells overexpressing pMIG-GFP control and pMIG-RBM3 **(B)** intracellular **(C)** extracellular (n=3). *, **, *** represents p-value < 0.05, 0.01 and 0.001 respectively.

**Supplementary Fig 9**

## S9 (A)

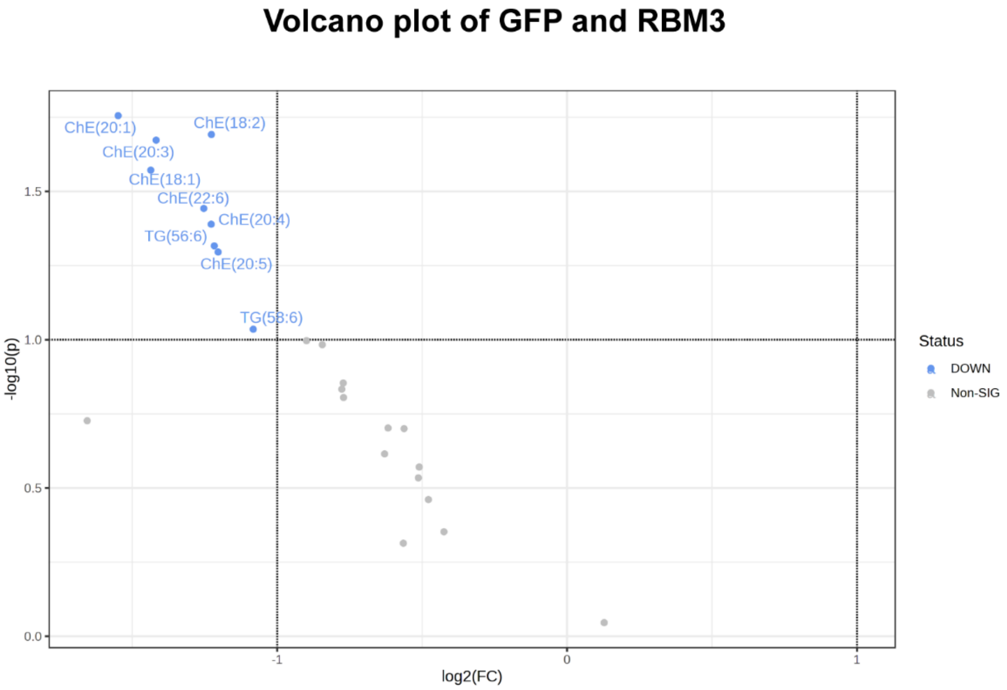


## S9 (B)

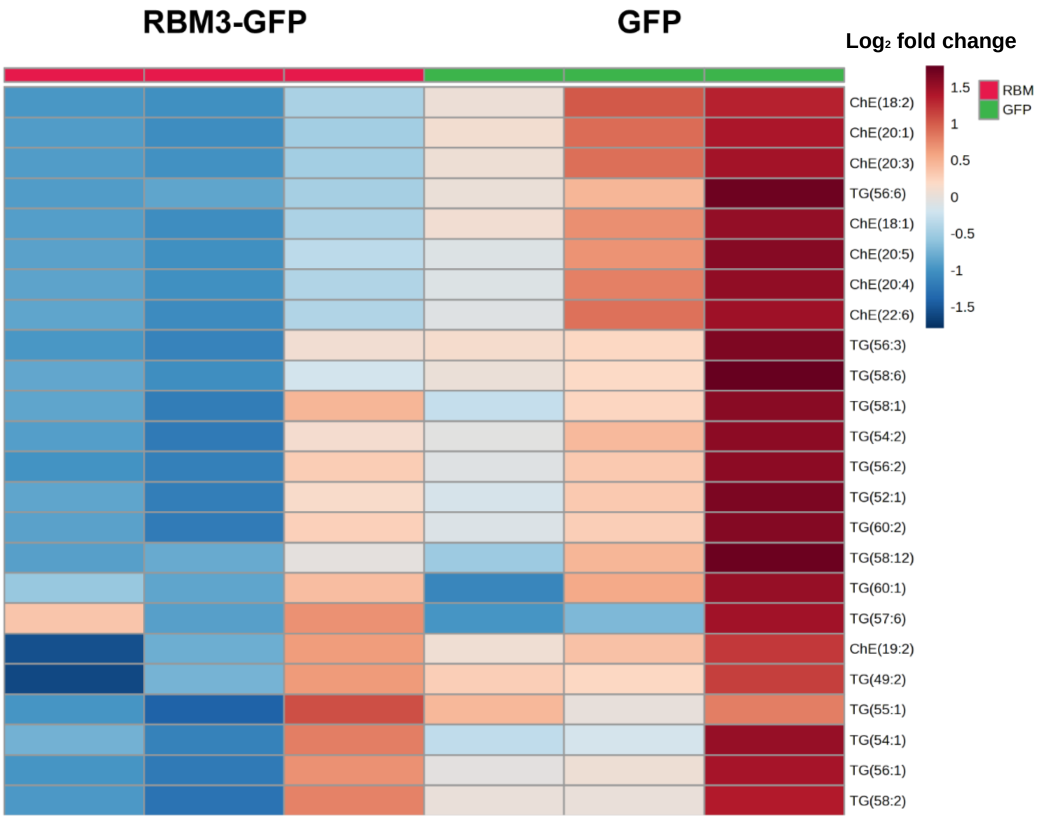

**Supplementary 9 (A)** Volcano plot of GFP and RBM3 overexpression where the x-axis represents the log of fold change and the y-axis represents -log of p-value . Grey dots represent non-significant lipid species and blue dot represents significantly low levels of lipid species (n=3). **(B)** Heat map showing levels of cholesterol esters and triglycerides using C2C12 cells overexpressing pMIG-GFP control and pMIG-RBM3 (n=3). *, **, *** represents p-value < 0.05, 0.01 and 0.001 respectively.

**Supplementary Fig 10**

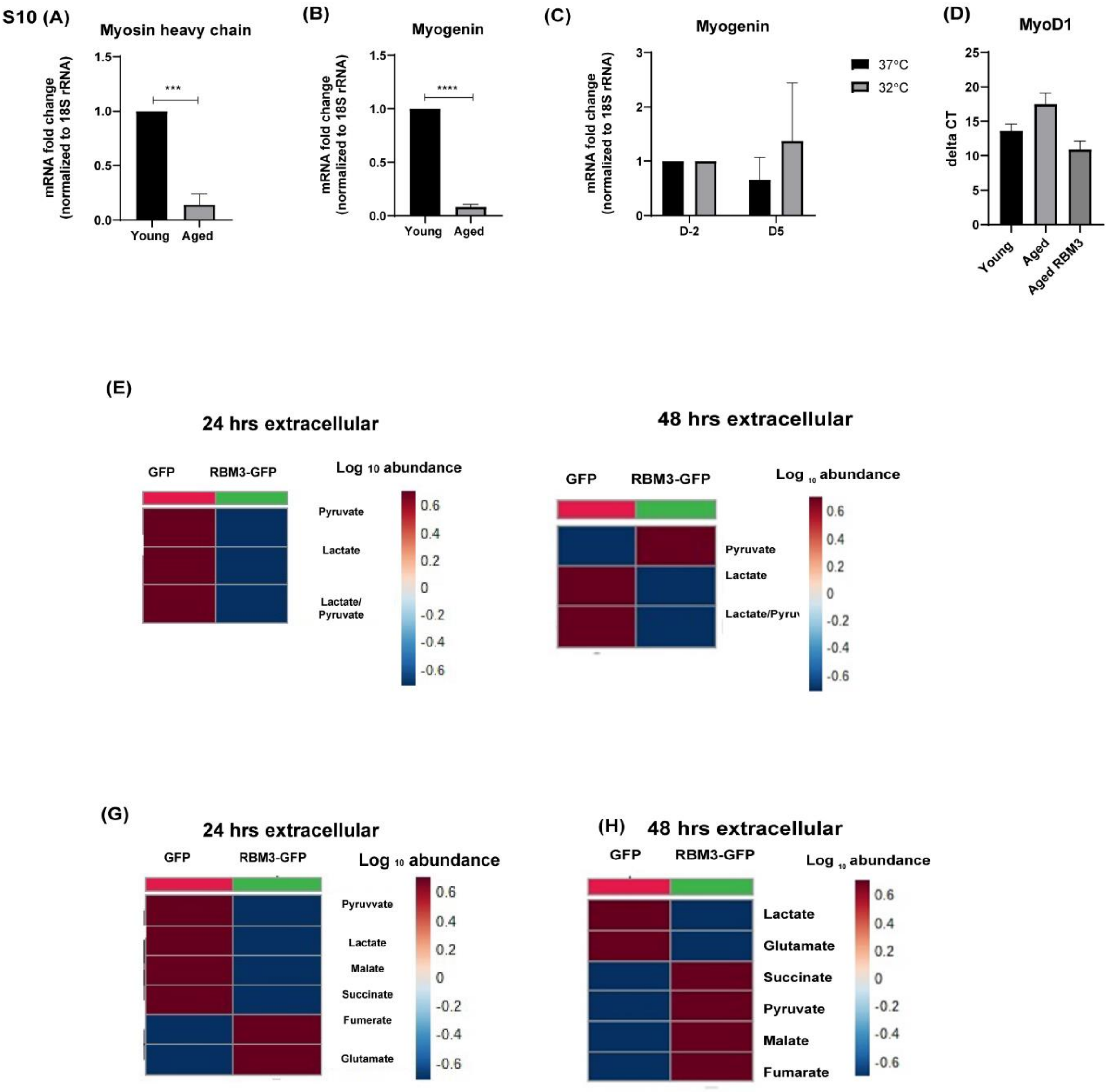

**Supplementary 10** mRNA expression levels of **(A)** *MyHC* and **(B)** *Myog* of aged and young myoblasts (n=3). **(C)** mRNA expression levels of *Myog* of aged myoblasts at $32^0$C compared to 37˚C (n=2). **(D)** Bar graph showing the comparative ΔCt values of young primary, aged primary and aged primary transfected with RBM3-GFP (n=2). Heat map showing levels of extracellular lactate/pyruvate ratio **(E)** 24 hrs. **(F)** 48 hrs. and TCA metabolites in aged mouse primary myoblasts transfected with RBM3-GFP and GFP control **(G)** 24 hrs. **(H)** 48 hrs. (n=3). *, **, *** represents p-value < 0.05, 0.01 and 0.001 respectively.

**Supplementary Fig 11**

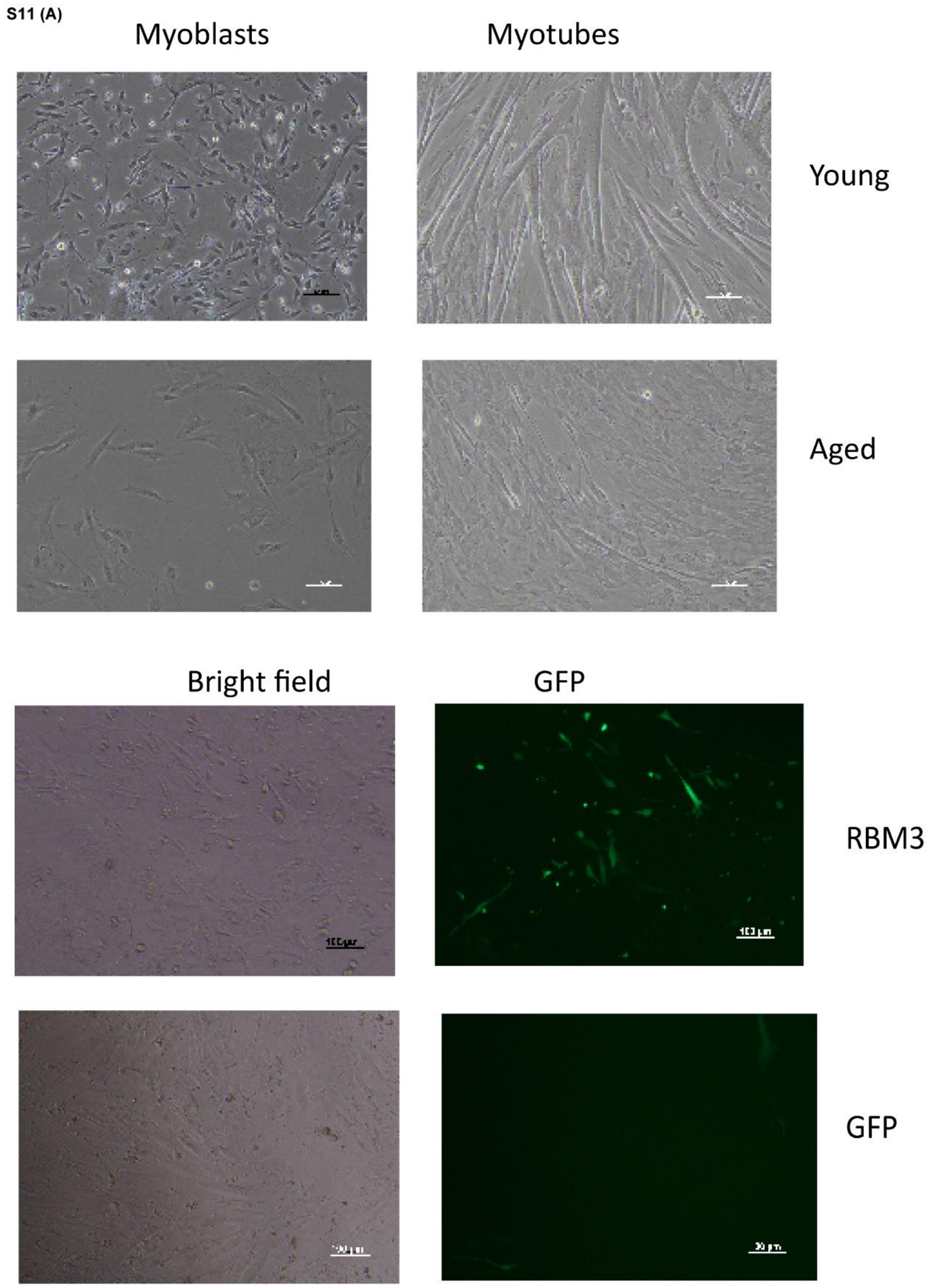


**Supplementary 11 (A)** Representative images of mouse primary myoblasts (young and aged) and differentiated cells (Scale bar, 100 um) **(B)** Representative images of mouse primary young myoblasts transfected with pMIG-RBM3 and pMIG-GFP in bright field and epifluorescence (Scale bar, 100 um).
